# Phage T3 overcomes the BREX defence through SAM cleavage and inhibition of SAM synthesis

**DOI:** 10.1101/2023.02.27.530186

**Authors:** Aleksandr Andriianov, Silvia Triguis, Alena Drobiazko, Nicolas Sierro, Nikolai V. Ivanov, Maria Selmer, Konstantin Severinov, Artem Isaev

## Abstract

Bacteriophage T3 encodes a SAMase that through cleavage of S-adenosyl-methionine (SAM) circumvents the SAM-dependent Type I Restriction-Modification defence of the host bacterium *Escherichia coli*. Here, we show that the SAMase also allows T3 to evade BREX defence. SAM degradation weakly affects BREX methylation of host DNA, but completely inhibits the defensive function of BREX, suggesting that SAM is required as a co-factor for BREX-mediated exclusion of phage DNA. The anti-BREX activity of the T3 SAMase is mediated by two independent mechanisms: enzymatic degradation of SAM and downregulation of SAM synthesis through direct inhibition of the host SAM synthase MetK. We determined a 2.8 Å cryo-EM structure of the eight-subunit T3 SAMase-MetK complex. Structure guided mutagenesis of the SAMase-MetK interface revealed that the interaction with MetK stabilizes the T3 SAMase *in vivo*, thus further stimulating its anti-BREX activity. This work provides insights in the versatility and intricacy of bacteriophage counter-defence mechanisms and highlights the role of SAM as an important co-factor of diverse phage-defence systems.

## Introduction

Evolutionary interactions between bacteriophages and their microbial hosts are often described in terms of an arms race. Prokaryotes developed a broad arsenal of measures to interfere with viral infection.^1^ The genomes of most free-living bacteria contain compact defence islands where genes encoding various defence systems are concentrated.^2,3^ Likewise, in mobile genetic elements, “anti-defence islands” containing genes responsible for the inhibition of host immunity, for instance, restriction-modification (R-M) or CRISPR-Cas systems,^4^ tend to form. Given that the diversity of known phage defence systems has significantly expanded in recent years,^5–8^ multiple host defence inhibitors should be encoded in phage genomes and mobile genetic elements. Indeed, phage proteins that target the recently recognized CBASS, Pycsar and Thoeris host defence systems have been identified.^9–11^ One can also expect that some phage anti-defence proteins could simultaneously counter unrelated defence systems. In line with this expectation, we showed that the product of *0*.*3* gene of the phage T7, a well-characterized inhibitor of Type I Restriction-Modification (R-M) systems,^12,13^ also inhibits the BREX defence of its host.

Here, we investigate the anti-BREX activity of the *0*.*3* gene product of a T7 relative - bacteriophage T3. Unlike T7, T3 inactivates host Type I R-M systems with a phage-encoded S-Adenosyl-L-Methionine (SAM) lyase (SAMase). The SAMase activity in lysates of T3-infected *E. coli* was documented as early as 1965, and it was proposed that T3 SAMase hydrolyses SAM to 5’-methylthioadenosine (MTA) and L-homoserine.^14^ However, recent work demonstrated that T3 SAMase instead cleaves SAM in a lyase reaction (Figure 1A).^15^ The T3 SAM lyase is encoded by the *0*.*3* gene, which is the first gene transcribed upon entry of the T3 genome into the cell. As a consequence, the SAM-cleaving activity is manifested at the earliest stage of phage infection.^16^ Mutations in the *0*.*3* gene make the phage unable to infect hosts carrying Type I R-M systems.^17^ A similar phenotype is observed for the T7 phage carrying mutations in its *0*.*3* gene, which encodes the Ocr protein. However, the anti-restriction mechanisms of the *0*.*3* gene products of the two phages are completely different. The T7 Ocr is a DNA-mimicking protein that binds the modification or restriction enzymes,^18,19^ while the T3 SAMase was proposed to inhibit the activity of R-M systems by depleting the pool of available SAM.^17^ Since Type I R-M systems, such as EcoKI, require SAM not only as a methyl donor for DNA modification but also as a cofactor for the DNA cleavage reaction,^20,21^ SAM depletion inhibits both activities. In the case of T3, a variant with a point mutation in the *0*.*3* gene was described that was seemingly devoid of SAMase activity and yet retained the ability to infect bacterial hosts carrying the EcoKI system.^17,22^ It was therefore proposed that the T3 SAMase could directly inhibit EcoKI, though no direct protein-protein interaction between the SAMase and the EcoKI system components was demonstrated.^23^ In addition, an unidentified non-EcoKI host protein was reported to copurify with the SAMase from T3-infected *E. coli* cultures.^16^ The role of this interaction, if any, and the detailed anti-restriction mechanism of the SAMase remains unknown.

**Figure 1.**
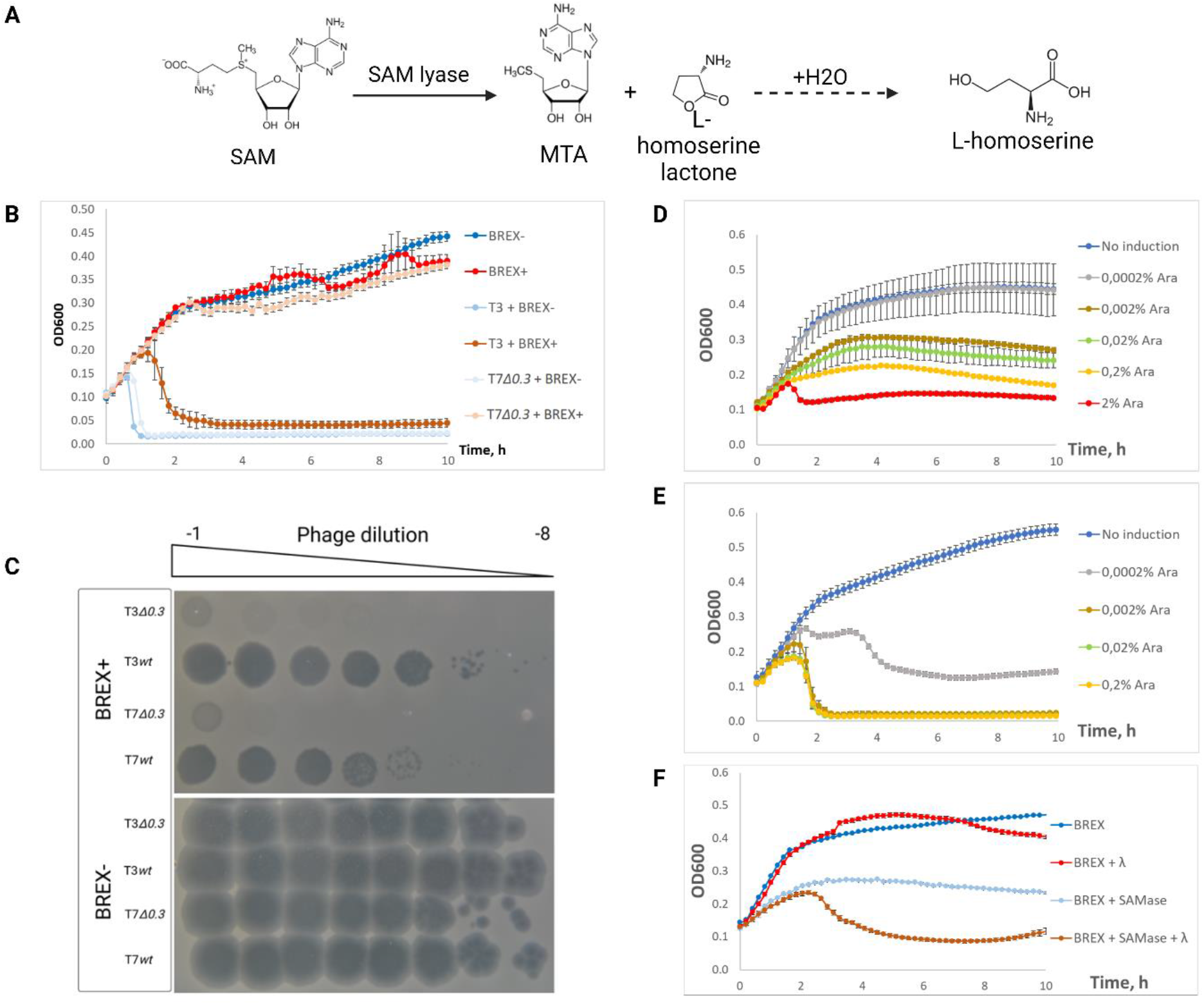
Phage T3 SAM lyase inhibits BREX defence. (A)Cleavage of S-Adenosyl-L-Methionine (SAM) into methyl-thioadenosine (MTA) and L-homoserine lactone catalysed by T3 SAM lyase. L-homoserine lactone is spontaneously hydrolysed to L-homoserine. (B)Growth curves of cultures infected with wild-type T3 or T7*Δ0*.*3* at MOI = 0.1. “BREX-” is *E. coli* BL-21 transformed with an empty pBTB-2 vector, “BREX+” cells are transformed with pBTB-2 derivative carrying *E. coli* HS BREX gene cluster transcribed from its native promoter. (C)Serial dilutions of T3 and T7 phages (wild-type or *Δ0*.*3* mutants) spotted on BREX- and BREX+ BL-21 *E. coli* lawns. Representative plates from an experiment performed in three biological replicates are shown. (D)Growth curves of BREX+ *E. coli* BW25113 culture transformed with compatible pBAD SAMase expression plasmid at different concentrations of L-arabinose. (E)Growth curves of BW25113 BREX+ culture carrying pBAD SAMase infected at MOI = 1 with phage T7*Δ0*.*3* in the presence of indicated concentrations of inducer. Phage was added at *t=0*, while SAMase expression was induced 30 minutes prior the infection. (F)Growth curves of BW25113 BREX+ culture carrying pBAD SAMase infected at MOI = 1 with phage λ_vir_ in the presence of 0.02% arabinose. Phage was added at *t=0*, while SAMase expression was induced 30 minutes prior the infection.

Development of the concept of defence islands – genomic loci enriched with immunity genes – promoted the discovery of multiple novel phage resistance systems.^2^ One group of such systems, which includes BREX and DISARM, encode methyltransferases and, similar to classical R-M systems, methylate host DNA to distinguish it from unmethylated invading DNA.^24–29^ However, these systems do not fit into the standard R-M classification, since they lack an obvious restriction endonuclease component but instead carry proteins not associated with characterized defence systems. BREX systems are found in about 10% of bacterial and archaeal genomes. The most widespread Type I BREX systems consist of six core components: BrxX (PglX) – a N6-methyladenosine methyltransferase; BrxZ (PglZ) – a putative alkaline phosphatase; BrxC – a putative ATPase; BrxL – a putative Lon-like protease, and two small proteins of unknown function (BrxA, BrxB). BREX prevents the accumulation of viral DNA in infected cell through a mechanism that remains to be identified. Known BREX methylation sites are asymmetric (GGTA**<u>A</u>**G in *E. coli* or TAGG**<u>A</u>**G in *Bacillus cereus*) and methylation occurs only at one DNA strand, as is also the case for Type ISP, IIL or III R-M systems whose recognition sites are also asymmetric.^30–33^ In Type ISP and Type III R-M systems self-DNA cleavage of unmethylated sites, which must appear upon modified DNA replication, is avoided by a requirement for an interaction between restriction complexes bound to non-methylated sites located on different DNA strands. In these systems, methylation, translocation, and restriction activities are combined in a single protein complex and recognition of a non-methylated site does not directly trigger DNA cleavage. These complexes are charged with SAM that is used as a methyl donor to modify unmethylated sites. The DNA cleavage reaction also requires SAM as a co-factor (albeit the role of SAM in Type III R-M is ambiguous).^20,21,34–36^ Thus, SAM depletion is an efficient strategy that can be used by phages to inhibit complex defence systems that require SAM for both target modification and target cleavage. Here, we show that the T3 SAMase inhibits BREX defence of the host and provide evidence that the active stage of BREX defence, whatever the actual mechanism, is also SAM-dependent.

## Results

### The *0*.*3* gene of bacteriophage T3 is required to overcome BREX protection of the host

To test whether T3 SAMase, like its functional analogue T7 Ocr, overcomes BREX protection, we infected cultures of *E. coli* BL-21 cells transformed with an empty pBTB vector (BREX-cells), or pBREX AL, a pBTB vector expressing *E. coli* HS Type I BREX (BREX+ cells), with wild-type T3. At a multiplicity of infection (MOI) of 0.1, wild-type T3 lysed the BREX-culture ~1 hour after infection, while lysis of the BREX+ culture took about 2 hours to occur (Figure 1B). As expected, the BREX+ culture was completely protected from infection with T7*Δ0*.*3* used as a control. A one-step growth experiment showed that although T3 progeny release by BREX+ cells was delayed compared to the BREX-control, the phage burst was not affected, consistent with the lack of efficient defence (Figure S1A,B).

We constructed a T3 mutant lacking the *0*.*3* gene (see Methods and reference^37^) and tested it for the ability to infect BREX+ cells. Similar to the wild-type T7, wild-type T3 produced plaques on a BREX+ lawn, though the titer was decreased by an order of magnitude and the plaques were smaller than those formed on the BREX-lawn. In contrast, T7 and T3 lacking their *0*.*3* genes did not form plaques on BREX+ lawns. We conclude that, similar to T7 gp0.3, gp0.3 of T3 is essential for overcoming the BREX defence of the host (Figure 1C).

### Expression of T3 SAMase is sufficient to shut off the BREX defence

SAM is an essential metabolite used in a variety of biosynthetic pathways either as a cofactor or a substrate. Therefore, SAM depletion is toxic.^38–41^ To overcome this, we cloned the T3 *0*.*3* gene into the pBAD L24 vector^12^ and expressed it from an arabinose-inducible promoter. Since two alternative start codons were predicted for the T3 SAMase ORF, resulting in 152 and 125 aa-long products, plasmids expressing both ORFs were constructed. Expression of the shorter ORF did not produce an anti-restriction effect (see below), therefore, only the plasmid expressing the longer, 152-codon ORF was used in further experiments. BW25113 *E. coli* BREX- or BREX+ cells were transformed with the T3 SAMase expression plasmid and their growth in the presence of L-arabinose inducer was monitored. In agreement with earlier data,^38^ induction with high (2%) concentrations of L-arabinose strongly inhibited growth; however, lowering the concentration of the inducer reduced or eliminated this toxicity (Figure 1D). In the absence of inducer, the BREX+ culture was protected from T7*Δ0*.*3* infection, while induction of T3 SAMase synthesis, with as low as 0.0002% L-arabinose, resulted in the lysis of infected cultures (Figure 1E). A similar effect was observed when the BREX+ culture expressing T3 SAMase was infected with λ_vir_ (Figure 1F), an unrelated phage that does not encode either homologs or analogs of T3 and T7 gp0.3 proteins. Therefore, we conclude that expression of T3 SAMase alone is sufficient to interfere with the antiviral defence by BREX.

To investigate whether SAM depletion could be a general strategy of BREX defence inhibition, we tested ORF1 and Svi3-3 SAM lyases, previously identified from bacteriophage metagenomic DNA libraries. Both of these proteins cleave SAM *in vitro*.^42^ The BREX defence was unaffected in cells expressing ORF1 and Svi3-3 from plasmids (Figure S1C,D). This could be due to the considerably lower SAM lyase activity of ORF1 and Svi3-3 compared to T3 SAMase.^15,42^ Alternatively, the result may indicate that the T3 SAMase counteracts BREX by means other than SAM cleavage.

### The anti-BREX activity of the T3 SAMase is not directly associated with inhibition of DNA methylation

In *E. coli*, Dam, Dcm, or R-M-specific methylation is suppressed upon T3 SAMase expression.^38,43^ To investigate whether the anti-BREX activity of T3 SAMase is a simple consequence of inhibition of BREX-specific methylation, we performed PacBio sequencing of genomic DNA from BREX+ cells expressing the SAMase (induction with 0.2% arabinose). The BW25113 strain used in this analysis has an EcoKI *r-m+* genotype, and thus, in addition to BREX methylation (GGTA**<u>A</u>**G), we were able to monitor the state of EcoKI (GC**<u>A</u>**CNNNNNNGTT and A**<u>A</u>**CNNNNNNGTGC) and Dam (G**<u>A</u>**TC) methylation (Figure 2A, S2, and Table S1). Sites with a QV value of >30 were considered as modified. While T3 SAMase expression resulted in ~2-fold reduction in the amount of Dam and EcoKI methylated sites, ~97% of BREX sites remained methylated. While this was not further investigated, the observed lack of inhibition of BREX methylation may be due to the higher abundance of plasmid-borne BREX methyltransferase compared to chromosomally-encoded EcoKI and Dam methyltransferases. At same conditions, induction of phage T7 gene *0*.*3* expression had a more profound effect, decreasing BREX methylation by ~20% and EcoKI methylation by ~60% (Figure 2A, S2).

**Figure 2.**
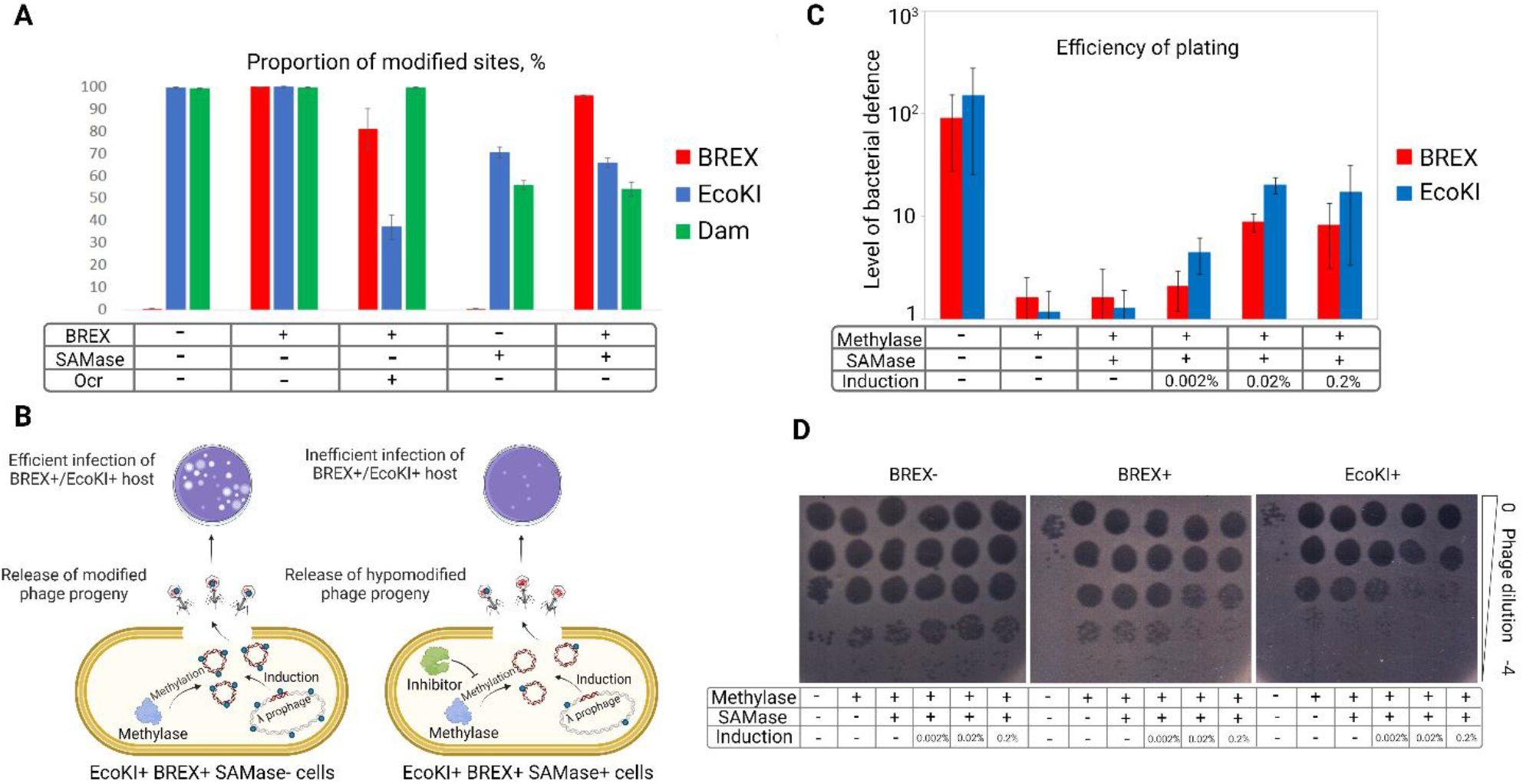
Anti-BREX activity of SAMase is not associated with inhibition of DNA methylation. (A)Proportion of modified Dam, EcoKI, and BREX sites in the genomic DNA of *E. coli* BW25113 expressing T3 SAMase or T7 Ocr in the presence of 0.2% arabinose as revealed by PacBio sequencing. (B)Phage λ_ts_ induction system for study of DNA methylation activity of the host. (C)Efficiency of plating of λ_ts_ phage induced from non-methylating (*ΔhsdM* EcoKI *m-r-* and BREX-) or methylating (EcoKI *m+r-* and BREX+) BW25113 *E. coli* hosts in the absence and in the presence of T3 SAMase. The level of defence is calculated as the ratio of phage titer obtained on the lawn of non-restricting (BW25113) relative to restricting (BREX+ or EcoKI+) hosts. AB1157 strain was used as an EcoKI+ (*m+r+*) restricting host. (D)Representative plates from a triplicate experiment shown in (C).

The effect of T3 SAMase on BREX methylation was also assessed using a λ_ts_ prophage induction assay,^25^ with EcoKI methylation of phage DNA monitored as an internal control. Fully methylated λ_ts_ phage induced from a EcoKI+/BREX+ host evades defence by both systems. Reduced methylation activity during prophage induction is expected to result in production of partially- or non-methylated λ_ts_ progeny that should be restricted on EcoKI+ or BREX+ lawns (Figure 2B). Phage λ_ts_ produced from the EcoKI+/BREX+ host at conditions of T3 SAMase induction with 0.2% arabinose had an intermediate (compared to fully-methylated or non-methylated variants) ability to propagate on BREX+ or EcoKI+ lawns, indicating that the EcoKI and BREX methylation activities in induced cells were compromised, though not completely inhibited (Figure 2C,D). Reducing the amount of arabinose to 0.002% resulted in the production of λts progeny that was fully protected from BREX, suggesting a complete or almost complete methylation of phage genomes. Yet, as shown above (Figure 1E), induction of SAMase synthesis with 0.002% arabinose was sufficient to inhibit BREX defence.

Together, the data presented above demonstrate that while T3 SAMase suppresses SAM-dependent methylation, its anti-BREX effect does not stem from the inhibition of methylation *per se*. Type I R-M systems rely on SAM for the assembly and/or activity of their restriction complexes,^20^ and SAM depletion directly deactivates restriction by these systems. The inhibition of BREX defence by T3 SAMase at conditions when BREX methylation was not affected, shows that the active stage of BREX defence, similar to Type I R-M systems, is also SAM-dependent. Induction of SAMase synthesis in cells harboring a Type II R-M system EcoRV, where restriction is SAM-independent and decoupled from methylation, leads to increased toxicity compared to the BW25113 control (Figure S1E,F). In contrast, presence of BREX+ or EcoKI+ systems does not increase SAMase toxicity (Figure S1E,F), further supporting that BREX defence is SAM-dependent.

### Catalytically deficient SAMase mutants retain partial anti-BREX activity

Multiple T3 phage mutants with compromised SAMase activity were obtained in the 1970-s. In most cases, the *in vitro* deficiency in SAM cleavage correlated with the lack of anti-restriction.^17^ However, one *0*.*3* mutant, 3356, lost the ability to degrade SAM but retained the anti-restriction activity.^17^ Further work on this mutant led to a hypothesis that the gp0.3 SAMase contributes to anti-restriction through mechanisms other than SAM degradation.^23^ Unfortunately, the 3356 mutant was lost, and the nature of the mutation remains unknown. We set out to reproduce the 3356 phenotype by testing catalytically deficient T3 SAMase mutants for anti-BREX activity. Based on the recently-published structure of the Svi3-3 SAMase^15^ and location of conserved active-site residues, we constructed T3 SAMase single-substitution mutants that were predicted to affect catalytic activity: E67Q, E68Q, Q94A, and D95A, as well as double mutants E67Q/E68Q and E68Q/Q94A (Figure S3A). BREX+ cells expressing these mutants from plasmids were checked for their ability to mount BREX defence against the T7*Δ0*.*3* infection (Figure 3A, S3B). E67Q, bearing a substitution of a non-conserved residue, behaved like wild-type. All other mutants showed reduced but clearly detectable anti-BREX activity.

**Figure 3.**
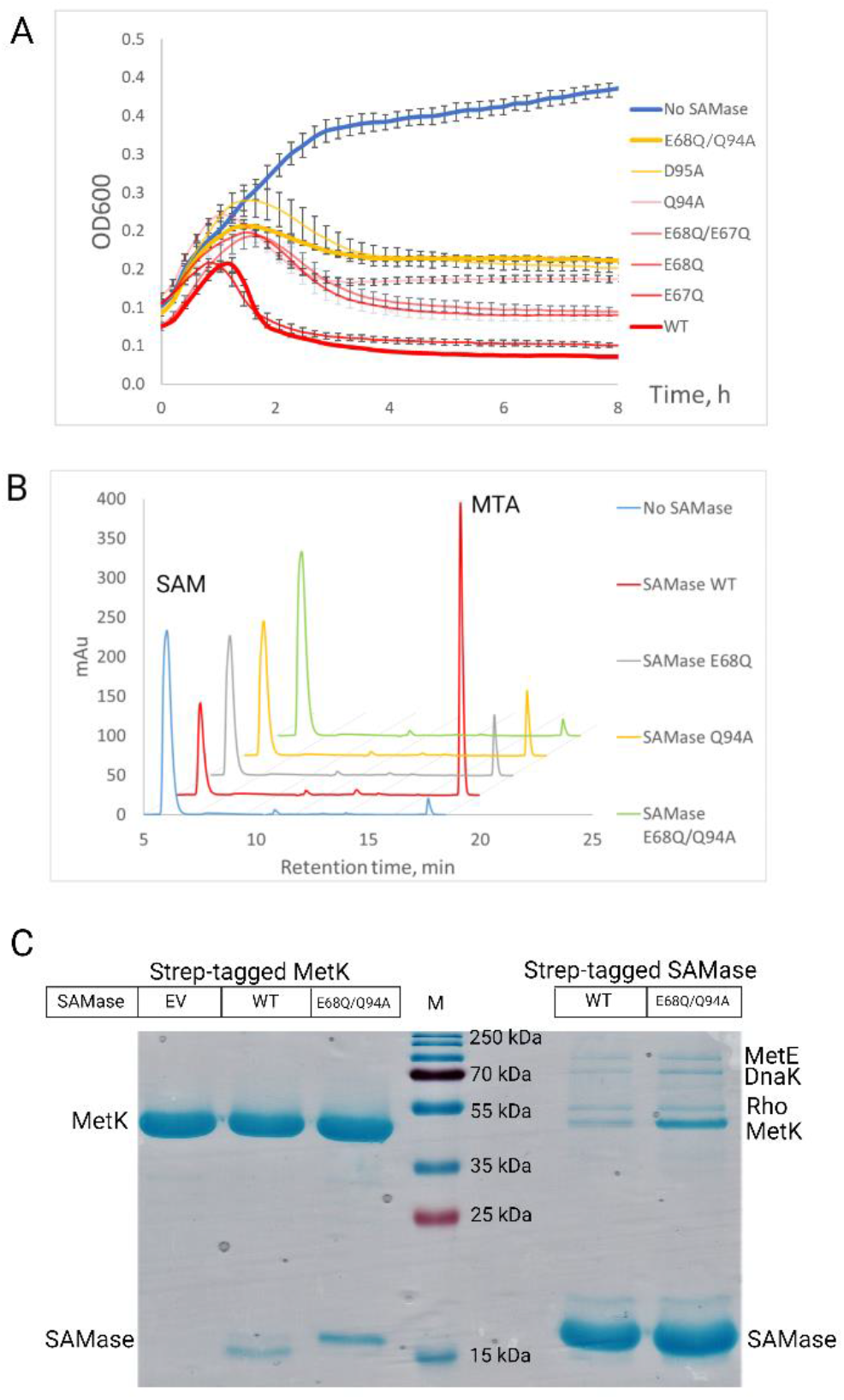
A catalytically-deficient SAMase mutant retains anti-BREX activity and interacts with SAM synthase MetK. (A)Growth curves of a BREX+ cultures producing different C-His SAMase mutants and infected with T7*Δ0*.*3* (MOI = 1, 0.2% arabinose). (B)Catalytic activity of wild-type, E68Q, Q94A, and E68Q/Q94A C-His SAMase mutants *in vitro*. HPLC separation of reaction substrate (SAM) and product (MTA) after 30 minute-incubation of 1 mM SAM with 500 nM of SAMase. Relative concentrations of substrate (SAM) and one of the products (MTA) were determined based on peak volume, and retention times were validated with SAM and MTA standards. No SAMase – incubation of 1 mM SAM without the addition of SAMase to estimate spontaneous SAM degradation. Absorbance was measured at 260 nm. Profiles are presented with an offset. (C)Right lanes - results of a pull-down from lysates of *E. coli* BW25113 expressing Strep-tagged SAMase from a pBAD vector (wild-type or E68Q/Q94A mutant). On the left - pull-down in conditions of Strep-tagged MetK expression from a pTG vector in the presence of non-tagged pBAD SAMase. A Coomassie stained 4-20% SDS gel is shown. The identity of all proteins was determined by MALDI-TOF mass spectrometry. EV – Empty Vector; M – Marker, PageRuler Prestained Plus.

We focused on the E68Q/Q94A double mutant and respective single mutants since the corresponding residues in Svi3-3 SAMase are critical for coordination of the reactive SAM conformation and a thus a defect in SAM binding is expected upon their substitution. Wild-type or mutated versions of recombinant T3 SAMase were purified, incubated with SAM for 30 minutes and the end-point reaction components were separated by reverse-phase HPLC. Compared to the wild-type, the amount of MTA generated from SAM by the E68Q and Q94A mutants was reduced 5-fold (Figure 3B). The E68Q/Q94A mutant did not produce detectable MTA at our conditions and was considered as catalytically inactive.

### T3 SAMase binds to the host SAM synthase MetK and inhibits SAM synthesis

Since the E68Q/Q94A mutant retained anti-BREX activity *in vivo*, it provided an entry point to study the earlier hypothesized additional anti-host defence function of T3 SAMase. Early reports indicated that a ~40 kDa host factor not related to R-M systems proteins copurifies with the T3 SAMase.^38^ Other work suggested direct inhibition of the EcoKI restriction complex by SAMase *in vitro*.^23^ Given this evidence, we monitored the interactions between the T3 SAMase and host proteins using a pull-down assay with a C-terminally Strep-tagged wild-type or E68Q/Q94A SAMase. No interaction with components of the BREX defence system in BREX+ cells extracts was detected (data not shown). However, at least three host proteins reproducibly copurified with the SAMase (Figure 3C). The most prominent co-purifying band contained MetK, the SAM synthase (or methionine adenosyltransferase; MAT) of *E. coli*. Given its size, 41.2 kDa, MetK is likely the host factor observed in the earlier work.^38^ The two weaker bands were identified as methionine synthase (MetE), the enzyme catalysing the penultimate reaction of SAM synthesis, and the transcription termination factor Rho. The three potential interaction partners of the wild-type enzyme also co-purified with the catalytically inactive E68Q/Q94A mutant. To verify the interaction of T3 SAMase with MetK, we performed a reverse pull-down assay with C-terminally Strep-tagged MetK and found that both wild-type and E68Q/Q94A untagged SAMase co-purified with MetK (Figure 3C).

The results presented above suggest that interactions with MetK (and, possibly, MetE and/or Rho) may be responsible for the residual anti-BREX activity exhibited by the catalytically inactive SAMase mutant. Specifically, if the binding of SAMase were to inhibit the activity of MetK and/or MetE, which catalyze the two final steps in the SAM synthesis pathway, then even a catalytically inactive SAMase could reduce intracellular SAM levels. While our work was in progress, this hypothesis was confirmed by an independent study,^44^ which showed that a T3 SAMase-MetK complex reconstructed *in vitro* was deficient in SAM synthesis; the SAMase activity was not compromised.^44^ However, this *in vitro* study did not address whether inhibition of SAM synthesis by the SAMase contributes to anti-restriction *in vivo*.

### Cryo-EM structure determination of the T3 SAMase – MetK complex

Wild-type or E68Q/Q94A T3 SAMase with a C-terminal hexahistidine tag was overexpressed and purified in complex with endogenous *E. coli* MetK. All wild-type T3 SAMase eluted in complex with MetK after size-exclusion chromatograph. The catalytically inactive mutant E68Q/Q94A was produced at higher level and eluted in two peaks corresponding to the MetK complex and free SAMase.

The wild-type T3 SAMase-MetK complex was subjected to structure determination using single-particle cryo-electron microscopy. (Table S2, Figure S4). *Ab initio* reconstruction without symmetry showed particles with size and shape consistent with a 4:4 complex composed of a central MetK tetramer with one T3 SAMase molecule bound to the distant end of each subunit. 3D classification showed that 46% of the particles had full occupancy of T3 SAMase, and only these were used for the high-resolution reconstruction. The best map for the full complex was obtained from non-uniform refinement with D2 symmetry (Figure 4A, GSFSC resolution 2.8 Å), while the best map for T3 SAMase was obtained after local refinement using symmetry expansion and a mask around the SAMase (Figure 4B, GSFSC resolution 3.0 Å). After real-space refinement of T3 SAMase and MetK in the respective best map, the complete 4:4 complex (Figure 4C) was fitted in the map from non-uniform refinement.

**Figure 4.**
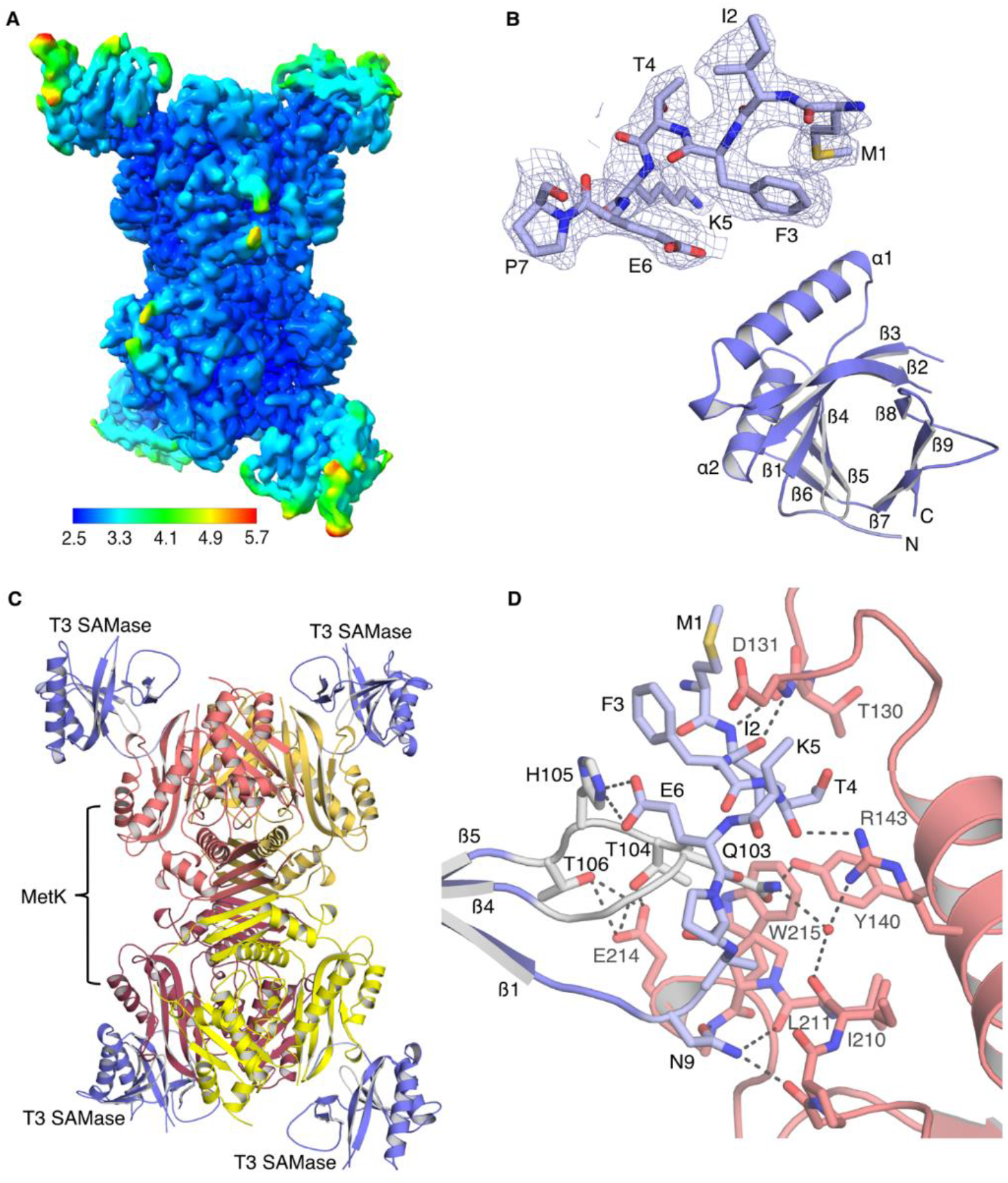
Cryo-EM structure of the T3 SAMase – MetK complex. (A)Cryo-EM map colored by resolution. (B)Map section after local refinement, overlaid with T3 SAMase (top), tertiary structure of T3 SAMase within the complex with MetK (bottom). (C)Structure of the T3 SAMase-MetK 4:4 complex. (D)Detailed interactions between T3 SAMase and MetK. Colors as in C with the N-terminal tail of the SAMase in light blue and the β4-β5 loop in light grey. SAMase-MetK hydrogen bonds involve the backbone of I2 and T4, and the side chain of N9 from the N-terminal tail and side chains of Q103, T104 and T106 from the β4-β5 loop. Q103 makes a direct hydrogen bond to Y140 in MetK and water-mediated hydrogen bonds to R143 and I210 in MetK. The N-terminal tail of T3 SAMase is stabilized by hydrogen bonds between H105 and E6. Hydrophobic contacts to T3 SAMase are mainly mediated by W215 and P212 in MetK but also between the aliphatic parts of T4 in the N-terminal tail and T130 in MetK.

### Structure of the T3 SAMase – MetK complex

The complex of T3 SAMase with MetK is composed of a central MetK tetramer with one T3 SAMase bound to each MetK subunit (Figure 4C). The MetK tetramer is formed by two tight MetK dimers, where each monomer consists of three similar alpha+beta domains, the N-terminal domain, the central domain and the C-terminal domain. Within the complex, T3 SAMase forms a monomeric nine-stranded beta barrel structure (β1-β9) with two helices (α1, α2) on the outside (Figure 4B, bottom). In the only previously known structure of phage-encoded SAM lyase, Svi3-3, the beta strands instead contribute to forming a trimer with active sites at the interface between the subunits.^15^ The T3 SAMase binds to MetK using a combination of hydrophobic contacts and hydrogen bonds involving side-chain and backbone atoms (Figure 4D, S5A,B). Residues M1, I2, F3, T4, K5, A8 and N9 from the N-terminal tail of T3 SAMase make contacts with the N-terminal domain and to the linker between the N-and C-terminal domains of MetK, while Q103, T104 and T106 from the β4-β5 loop in T3 SAMase form a network of hydrogen bonds with the N-terminal domain of MetK.

Analysis of SAMase sequences in the clustered non-redundant NCBI database with at least 30% sequence identity to the T3 enzyme shows that the MetK-interacting residues T4, N9, Q103 and T104, and H105 that stabilises the N-terminal tail, are strictly conserved (Figure S5C). This suggests that the ability to form a complex with MetK of the host bacterium is conserved among SAMases of phages infecting closely related hosts but not among more divergent SAMases, as evident by the lack of MetK-binding residues in Svi3-3 and ORF1 (Figure S3A).

### Stoichiometry and solution structure of the T3 SAMase-MetK complex

In a recent publication, a lower resolution polymeric structure of the T3 SAMase-MetK complex, linked together by SAMase dimers was presented.^44^ Our cryo-EM data show no sign of such polymers, which were proposed to be important for the physiological mechanism of the T3 SAMase. For this reason, we decided to confirm the stoichiometry of our complex using solution methods. To this end, molecular weight determination was performed using mass photometry with the E68Q/Q94A T3 SAMase-MetK complex (Figure 5A). A molecular weight of 239 kDa agrees perfectly with the calculated molecular weight for the 4:4 stoichiometry observed in the cryo-EM structure.

**Figure 5.**
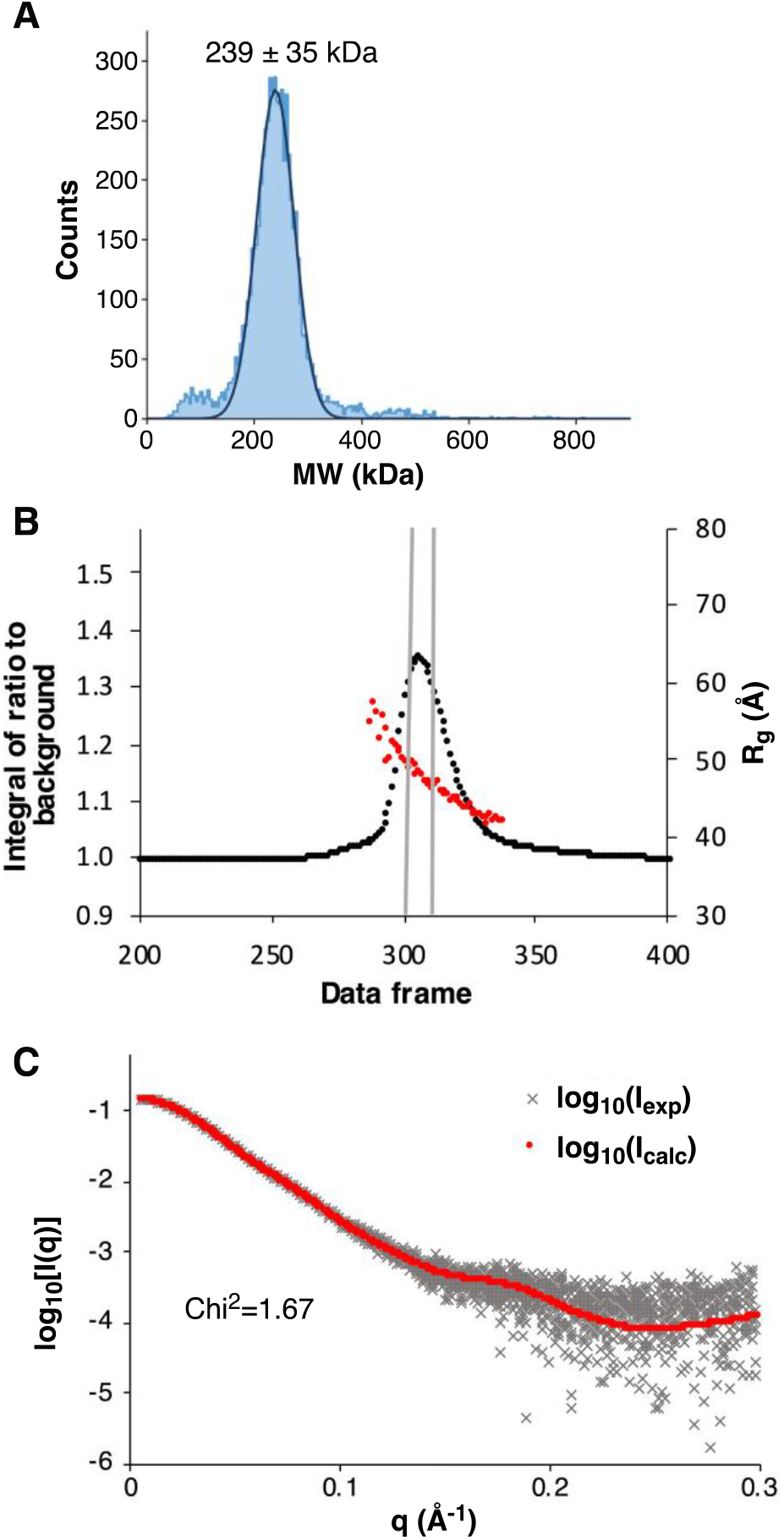
The T3 SAMase-MetK complex has a 4:4 stoichiometry in solution. (A)Mass photometry mass distribution histogram for the E68Q/Q94A T3 SAMase-MetK complex. The fitted Gaussian distribution accounts for 5635 counts (88% of total counts). (B)SEC-SAXS scattering intensity plot for the wild-type T3 SAMase-MetK complex. Red markers indicate the radii of gyration (R_g_) calculated from the individual scattering curves. Grey bars indicate the data frames used for further analysis. (C)Overlay of experimental scattering data (black) and the calculated scattering curve based on the 4:4 wild-type T3 SAMase-MetK cryo-EM complex structure (red). Equivalent plots for the E68A/Q94A T3 SAMase-MetK complex are presented in Figure S6.

Additionally, the wild-type and E68Q/Q94A T3 SAMase-MetK complexes were subjected to size-exclusion chromatography coupled to small-angle X-ray scattering (SEC-SAXS). The scattering intensity plots show no signs of larger oligomers or aggregates (Figure 5B, S6A), and the scattering curve shows excellent agreement with the size and shape of the complex in the 4:4 cryo-EM complex structure (Figure 5C, S6B). In conclusion, there is no indication of polymer formation in natively purified T3 SAMase-MetK complex and the stoichiometry in solution agrees with the single-particle cryo-EM reconstruction.

### Interaction with MetK stabilizes T3 SAMase *in vivo* and contributes to the anti-BREX activity

Based on the observed specific contacts between the N-terminal tail and the β4-β5 loop of T3 SAMase with MetK (Figure 4D), we designed a series of mutations in these regions to disrupt the binding. To prevent the interaction with the N terminus, we created 5- and 10-aminoacid N-terminal truncations of the SAMase and introduced mutations of three amino acids predicted to be crucial for the interaction (I2A/T4S/K5S). Mutations were introduced in both wild-type and E68Q/Q94A plasmid-borne hexahistidine-tagged SAMase. The anti-BREX activity of mutants was tested by monitoring the ability of BREX+ cells expressing SAMase variants to withstand phage infection, while the binding to MetK was tested using a pull-down assay from cell extracts (Figure 6A). The Δ5, Δ10 and I2A/T4S/K5S variants showed reduced or zero anti-BREX activity in the wild-type SAMase background; abolished BREX inhibition in the background of the catalytically dead E68Q/Q94A SAMase and did not co-purify with MetK (Figure S7A-C). However, the yield of purified proteins was strongly reduced compared to the wild-type or E68Q/Q94A SAMase, suggesting potential folding problems (Figure 6B). To overcome this, we introduced milder mutations in the SAMase N-terminus (ΔI2, which is equivalent to ΔM1/I2M, and T4A) and in the β4-β5 loop (Q103A and T104A) (Figure 6A). The T4A single substitution mutant was produced at the highest level (but still below the wild-type), (Figure 6B) and was purified in good yield. To confirm that the mutant was correctly folded, we performed a SAMase end-point assay and found that T4A was fully active (Figure 6C). Yet, the anti-BREX activity of T4A was compromised in the wild-type background (Figure 6D), and was abolished in the catalytically-deficient E68Q/Q94A background (Figure 6E, left). Compared to the wild-type, the amount of MetK co-purified with the T4A SAMase was reduced. Moreover, the resultant complex was unstable as it readily dissociated during gel-filtration (Figure 6F, right).

**Figure 6.**
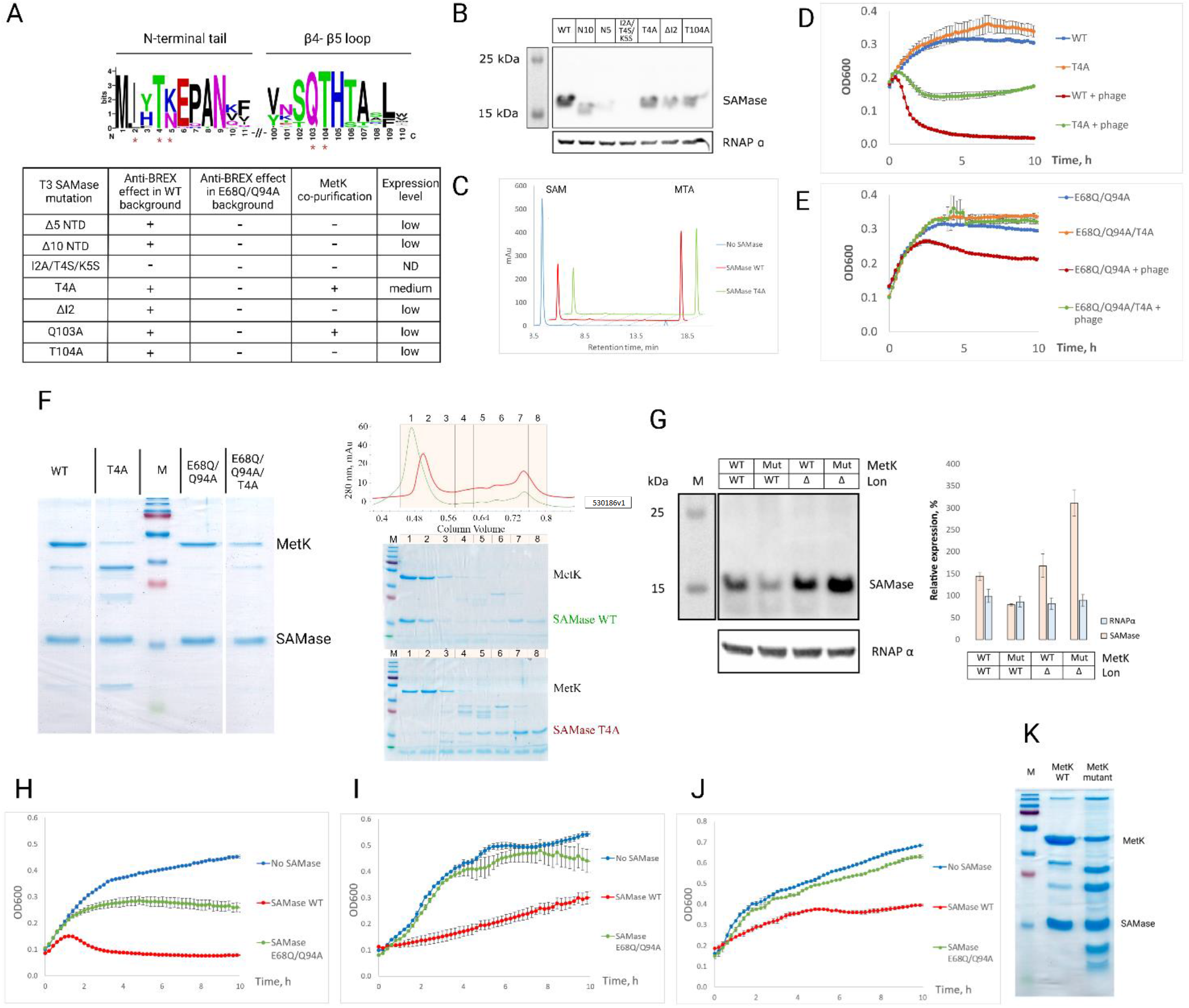
Interaction with MetK stabilizes T3 SAMase *in vivo* and contributes to its anti-BREX activity. (A)Mutations introduced into the MetK-interacting interface of T3 SAMase and their effects on anti-BREX activity and protein stability. Red stars on the logo indicate positions where amino acid substitutions were introduced. Expression level was estimated by western blot. *(B)In vivo* expression levels of wild-type or mutant His-tagged SAMase proteins as determined by western blot with anti-His antibodies. *E. coli* RNA polymerase subunit was used as a reference. (C)Activity of wild-type or T4A mutant T3 SAMase *in vitro*. HPLC separation of reaction substrate (SAM) and product (MTA) after 30 minutes’ incubation of 1 mM SAM with 500 nM of SAMase. No SAMase – incubation of 1 mM SAM without the addition of SAMase to estimate spontaneous SAM decay. Profiles are presented with an offset. (D-E) Growth curves of BREX+ cultures producing wild-type C-His SAMase (D) or E68Q/Q94A mutant with or without additional T4A mutation (E) upon infection with T7*Δ0*.*3* (MOI = 0.1, 0.2% arabinose). (F)A 16% SDS gel showing the results of pull-downs from lysates of *E. coli* BW25113 expressing His-tagged wild-type SAMase or indicated mutants. Protein concentrations were adjusted to achieve equal amount of SAMase in each lane. Wild-type and T4A SAMase eluates were further separated on a gel filtration column (right, top) and fractions were analyzed by SDS-PAGE (right, bottom). The identity of all SAMase and MetK bands was confirmed using MALDI-TOF MS analysis. M – Marker, PageRuler Prestained Plus. (G)A western blot showing the amounts of plasmid-borne His-tagged wild-type T3 SAMase in *E. coli* MG1655 strains carrying wild-type or *E. coli-to-N. gonorrhoeae* chimeric *metK* gene on the *lon*^*+*^ or *lon*^*-*^ background were determined by western blot. A signal from an RNA polymerase subunit was used for normalization (taken for 100%). Western blots were performed in triplicate, average band intensities were calculated in ImageJ and are presented on the right. (H-J) T3 SAMase has reduced anti-BREX activity in *E. coli* background with alternate MetK variants. Growth of *E. coli* MG1655 (H), MG1655 with a substitution of chromosomal *metK* for a homologue from *U. urealyticum* and additional *U. urealyticum metK* expressed from the pFlag vector (I), or MG1655 carrying several *E. coli-to-N. gonorrhoeae* mutations in its chromosomal *metK* (J) upon infection with phage T7*Δ0*.*3* (MOI = 1). Cells were BREX+ (carried the pBREX AL plasmid) and also harbored wild-type or E68Q/Q94A SAMase expression plasmid (induced with 0.2% arabinose). (K) A 16% SDS gel showing the results of pull-downs from lysates of *E. coli* MG1655 encoding wild-type (WT) or *E. coli-to-N. gonorrhoeae* (“mutant”) MetK and expressing His-tagged T3 SAMase. Protein concentrations were adjusted to load an equal amount of T3 SAMase in each lane. The identity of SAMase and MetK bands was confirmed using MALDI-TOF MS analysis. M – Marker, PageRuler Prestained Plus.

The combined observations that the T4A substitution decreased the SAMase:MetK complex stability and interfered with the anti-BREX function, while maintaining SAM cleavage, suggests that both MetK binding *and* SAMase activity are jointly required for full inhibition of BREX defence. To provide independent support for this conclusion, we decided to also test the effect of disruption of the SAMase-binding interface of MetK. The T3 SAMase was previously shown to interact with *E. coli* MetK but not with MetK from *Neisseria gonorrhoeae* or *Ureaplasma urealyticum*.^44^ Indeed, while expression of the wild-type SAMase resulted in a partial anti-BREX effect in the *E. coli* strain carrying the *U. urealyticum metK* instead of the *E. coli* homologue, presumably due to its SAM-degrading activity, the E68Q/Q94A mutant had no anti-BREX effect in this background (Figure 6H,I). Since *E. coli* carrying the *U. urealyticum metK* showed reduced viability, we additionally tested an *E. coli* MetK mutant with substitutions in the T3 SAMase interaction surface that introduced residues present in the *N. gonorrhoeae* MetK (D132P, V133T, I178V, I211V, A214P, W26L). The resulting hybrid MetK protein retains full catalytic activity but is not subject to inhibition by T3 SAMase *in vitro*.^44^ In this background, the toxicity of the wild-type SAMase was reduced, while the E68Q/Q94A mutant did not inhibit cell growth (Figure S7D,E). Importantly, the anti-BREX activities of wild-type and E68Q/Q94A SAMase were also severely reduced in this strain, confirming the importance of MetK binding for complete inhibition of BREX (Figure 6J). Pull-down experiments with His-tagged SAMase confirmed decreased efficiency of MetK co-purification with the MetK chimera (Figure 6K). Overall, we conclude that binding to MetK is required for full anti-BREX activity of T3 SAMase.

Unexpectedly, the wild-type SAMase was poorly produced in the strain with the *E. coli*-to-*N. gonorrhoeae metK* chimeric gene, mimicking the situation observed with SAMase variants carrying mutations in the MetK-binding surface (Figure 6G). This result suggests that disruption of the MetK:SAMase interface leads to decreased stability of the SAMase *in vivo*, regardless of which component of the MetK-SAMase complex is mutated. A possible explanation is that MetK protects SAMase from proteolytic cleavage. To test this hypothesis, we monitored the amounts of SAMase produced in strains with deletions of the common toxin-degrading proteases Lon or ClpXP.^45^ Indeed, deletion of the Lon protease gene resulted in increased production of wild-type SAMase and enhanced toxicity (Figure 6G, S7F). Yet, we were unable to confirm cleavage of SAMase or SAMase:MetK complexes by Lon *in vitro* (data not shown). Further studies are needed to decipher the role of Lon or other proteases in T3 SAMase degradation.

## Discussion

In this work we show that phage T3 possesses a powerful tool to inhibit both Type I R-M and BREX defences of the host. While the finding mirrors our earlier observation with phage T7, a relative of T3, the two phages rely on completely different mechanisms to achieve the same goal. T7 encodes Ocr, a DNA mimic that prevents the DNA binding by Type I R-M and BREX systems machinery, thus abolishing the defence by preventing viral genome recognition. T3 encodes a SAMase that degrades SAM, a universal donor of methyl groups. A decrease of intracellular SAM concentration should result in decreased levels of host DNA methylation. For simple defence mechanisms such as Type II R-M systems, the absence of methylation-mediated protection (compared to the unmodified phage DNA) shall cause destruction of host DNA by endonucleases. In the case of Type I R-M systems, SAM is required for both methylation and restriction activity^20^ and, therefore, at lowered SAM concentration, the chances of destruction of unmodified DNA decrease, benefiting the phage. While the mechanism of BREX-mediated exclusion is unknown, our data show that it also must be SAM-dependent.

We unexpectedly found that a catalytically dead E68Q/Q94A SAMase mutant is able to partially supress BREX defence. The residual anti-BREX activity is likely due to the retained binding of the SAMase mutant to MetK, the SAM synthase of the host. This binding inhibits SAM synthesis *in vitro*^44^ and, presumably, *in vivo*. In addition, SAMase becomes stabilized when bound to MetK *in vivo*, which should further decrease SAM levels. Also, binding of the SAMase to methionine synthase MetE may contribute to residual activity of the E68Q/Q94A mutant. The three-pronged model of T3 SAMase action is shown in Figure 7. The relative contributions of the two consequences of the SAMase-MetK interaction (inhibition of SAM synthesis by MetK and stabilization of the SAMase) to anti-BREX activity remain to be determined. In any case, this holistic mechanism of T3 SAMase action allows to quickly lower the intracellular SAM pool at the earliest stages of infection and consequently switch off the host defence.

**Figure 7.**
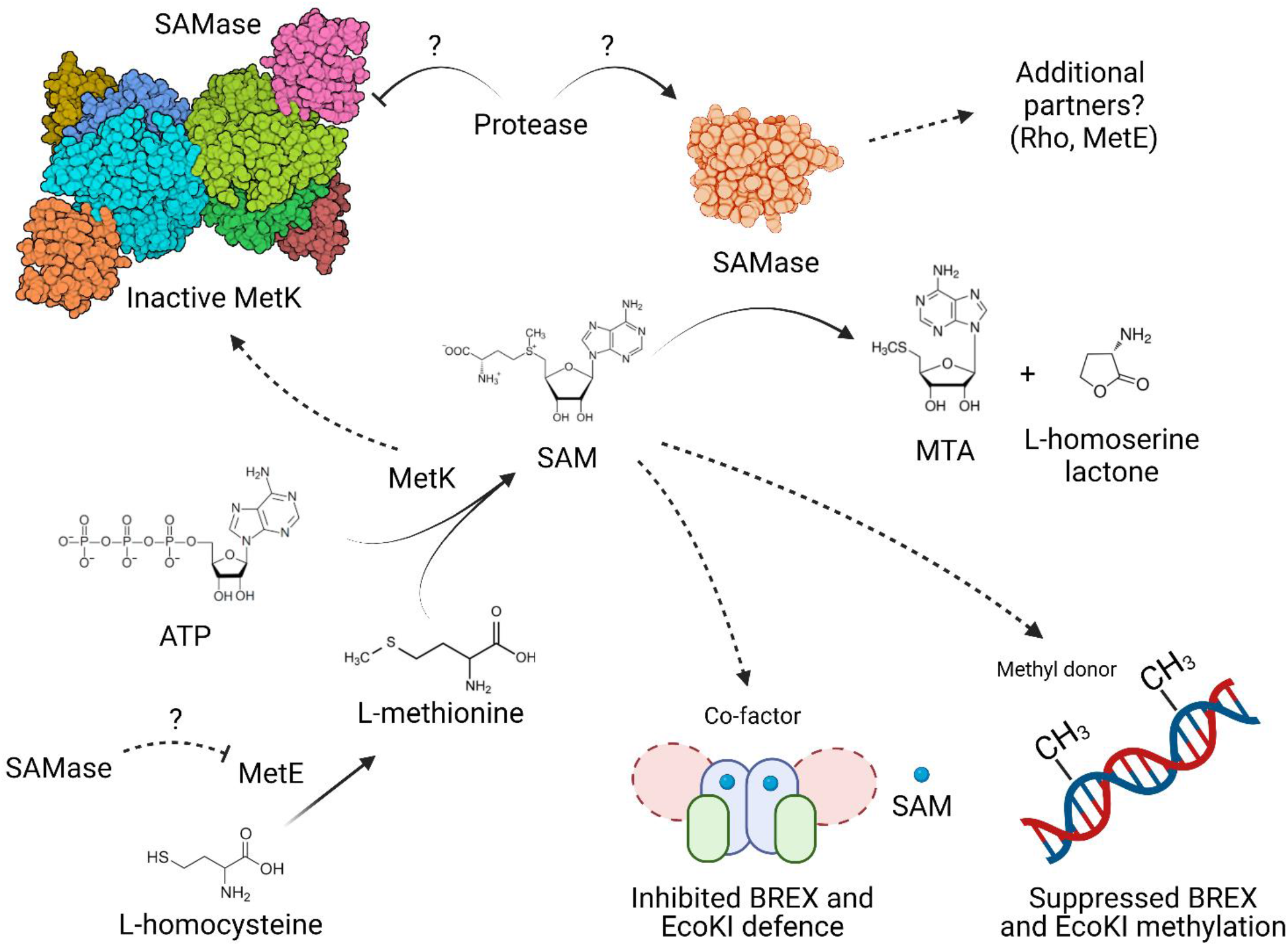
A model of T3 SAMase-mediated inhibition of BREX and EcoKI defence systems.

There is an apparent paradox in the observation that expression of T3 SAMase from a plasmid has virtually no (in the case of BREX) or partial (in the case of Type I R-M system) effect on host DNA modification while substantially affecting host defence. We hypothesize that during normal growth of uninfected cells, a lowered concentration of active (i.e., SAM-bound) modification enzymes is sufficient to maintain close-to-complete levels of host DNA methylation for both defence systems.

However, in cells expressing the T3 SAMase (from a plasmid or during T3 infection), the decrease in concentration of active, SAM-bound, restriction or exclusion enzymes provides a time-window where phages can take over the host metabolism and eventually lyse the cells, before being attacked by SAM-dependent complexes. The proposed dynamic interplay of methylation and restriction/exclusion activities underscores the similarities between the Type I R-M and BREX defences. In the case of BREX, the role of SAM may be direct as in the Type I R-M systems (i.e., as required co-factor for the exclusion reaction), or indirect (i.e., promoting increased affinity of BREX components to unmodified phage DNA).

Why has T3 phage developed an additional strategy to modulate intracellular SAM concentration, besides enzymatic cleavage? In *E. coli*, SAM acts as a co-repressor of the *met* regulon from which the enzymes involved in methionine biosynthesis are expressed. The feedback inhibition of methionine biosynthesis based on SAM concentration means that SAM degradation leads to up-regulation of the *met* regulon,^41,42^ counteracting the effect of SAM degradation. Since MetK converts methionine to SAM, its inhibition by direct interaction with T3 SAMase provides a more efficient way of lowering SAM levels.

The homogeneous, symmetric, T3 SAMase-MetK complex containing equimolar amounts (4:4) of both proteins observed here was prepared by overexpression of T3 SAMase in *E. coli*, and is consistent with the intracellular amounts of T3 SAMase (~10 000 molecules per cell early in phage infection^46^) and MetK (around 7000 in uninfected cells^47^). Results from gel filtration, mass photometry and SEC-SAXS experiments support the 4:4 stoichiometry, suggesting that the T3 SAMase has higher affinity to MetK than to itself. Thus, T3 SAMase monomers should bind to MetK tetramers until the latter are saturated. Based on the observation that monomeric T3 SAMase binds to MetK (this study), while free T3 SAMase purifies as a dimer,^44^ the native dimer interface may overlap with the MetK contact area. While our work was in progress, a polymeric T3 SAMase-MetK structure obtained from refolded T3 SAMase and separately purified MetK was reported.^44^ The complex polymers are held together by an asymmetric T3 SAMase dimer, possibly a refolding artefact. Our higher-resolution structure allowed unambiguous modelling of the SAMase-MetK interface, and the defined 4:4 oligomer is more likely to represent the complex formed during phage infection.

It is puzzling how binding of T3 SAMase 25 Å from the closest catalytic site of the MetK tetramer can cause inhibition. Simulations based on the polymeric T3 SAMase-MetK structure suggested that allosteric inhibition of MetK could be mediated through increased dynamics of the substrate-binding pocket.^44^ Even though the structure presented here is at significantly higher resolution (2.8 Å) than the previous, polymeric, T3 SAMase-MetK structure (3.6 Å), it is still difficult to draw conclusions from comparison with the previous crystal structures of the free MetK due to crystal-packing effects. The location of the T3 SAMase binding surface at the edge of the N-terminal domain and at the linker between the N-and C-terminal domains of MetK suggest that inhibition might be mediated by subtle effects on the domain and/or subunit arrangement of MetK. Further structural and biochemical studies are needed to clarify this mechanism of inhibition.

Since SAM-dependent restriction systems are widespread in the bacterial genomes and put a lot of pressure on phages, the primary role of phage-encoded SAMases, of which currently several hundred can be found, is likely the inhibition of host defences. However, targeting of SAM, an essential metabolite that is involved in a plethora of biosynthetic pathways, opens additional possibilities for the phage to manipulate the metabolism of the host. For example, the T3 phage when infecting starved *E. coli* cultures is able to enter a so-called “pseudolysogenic” state and persist in infected cell without replication.^48,49^ The ability to enter the pseudolysogenic state depends on the presence of the SAMase gene. Curiously, SAMase expression is required during a round of infection that precedes the infection of starved cells.^49^ Although the mechanism of pseudolysogeny is unknown, it might be expected that the T3 genome should be demethylated to enter the pseudolysogenic state. SAM is a precursor of quorum sensing (QS) signalling molecule autoinducer-2 (AI-2). Expression of the T3 SAMase was shown to decrease AI-2 production.^50^ In laboratory strains of *E. coli*, the AI-2 dependent regulation of gene expression is limited and accordingly AI-2 plays a negligible role in phage infection, although it influences the ability to form biofilms.^51–53^ However, in other bacteria, such as *Pseudomonas* and *Serratia*, QS signalling heavily affects phage infection through modulation of surface receptors, biofilm formation, and direct upregulation of expression of defence systems, such as CRISPR-Cas.^54–58^ Thus, quenching of the QS signalling should be beneficial for phages, and phage-encoded modulators of the QS pathways were indeed recently reported.^59,60^ Homologs of T3 SAMase are found in some phages infecting *Serratia* and *Pseudomonas*, and it would be of interest to determine whether these enzymes suppress QS signalling of the host. SAM is also required for the synthesis of polyamines, such as spermidine and spermine.^61^ Recent work demonstrated that polyamines inhibit phage replication and once released from lysed infected cells serve as signalling molecules to activate phage defence in neighbouring uninfected cells through the Gac/Rsm pathway.^62^ Again, it might be expected that phage-encoded SAMases would reduce the synthesis of polyamines and thus either protect the phage from direct inhibition or prevent the activation of host defences.

In conclusion, T3 SAMase represents an unusual example of a small multifunctional protein that makes use of several distinct mechanisms to lower the SAM concentration during infection. Besides its ability to inhibit BREX and Type I R-M defence systems, the T3 SAMase and its homologs have potential to manipulate additional SAM-dependent host processes leading to increased efficiency of phage infection.

## Methods

### Bacterial strains, plasmids, phages and growth conditions

The full list of bacterial strains, phages and plasmids used in this work can be found in Table S3. *E. coli* BW25113 was routinely used for experiments with pBAD T3 SAMase. BL-21 strain was used for the infection experiments with T3 phage. AB1157 was used for the experiments with EcoKI Type I R-M system. XL1-Blue strain was used for the transformation after molecular cloning. *E. coli* MG1655 was used as a background for the study of *metK* mutations, *E*.*coli-to-U*.*urealyticum metK* and *E*.*coli- to-N*.*gonorrhoeae metK* mutants^44^ were kindly provided by Dr. Shimon Bershtein. Unless otherwise indicated, cells were grown at 37°C in standard LB media with appropriate antibiotics. For the infection with phage λvir cells were supplemented with 0.2% maltose and 5 mM MgSO_4_. Phages were routinely propagated on BW25113 *ΔhsdM* (KEIO) to avoid EcoKI-specific methylation. BW25113 transformed with an empty vector pBTB-2 is referred to as BREX-through the text, while BREX+ represents BW25113 transformed with pBREX AL carrying Type I BREX 6 genes cluster from *E. coli* HS cloned with native promoters.^25^

### Plasmid construction

Primers used to generate plasmids used in this work are listed in Table S4. pBAD T3 SAMase was constructed by cloning 378-bp *0*.*3* gene from T3 genome into pBAD L24^12^ under control of *araBAD* promoter. PCR product was assembled with EcoRI-linearized pBAD L24 using NEBuilder HiFi DNA Assemly Master Mix (NEB). Strep-Tag II and His-tag C-terminally modified SAMase coding plasmids were constructed in a similar manner. MetK C-strep was cloned under control of *araBAD* promoter into pTG^25^ using BW25113 genomic DNA as a source of *metK* gene and Gibson Assembly with SacI-linearized vector. All mutations in pBAD T3 SAMase were introduced using Q5 Site-Directed Mutagenesis Kit or KLD Master Mix (NEB). Plasmids were transformed into chemically competent XL1-Blue cells (Evrogen). Phusion DNA-Polymerase (NEB) was used for the generation of PCR products for cloning, while Taq 5x ScreenMix (Evrogen) was used for the screening of colonies and routine PCR. Identity of all constructs was verified by Sanger sequencing (Evrogen).

### Bacterial culture growth dynamics

Growth of bacterial cultures to study phage infection or toxicity of SAMase was carried at 37°C in a 96-well format using EnSpire Multimode Plate Reader (Perkin Elmer). Overnight bacterial cultures were diluted 100-fold in LB medium with appropriate antibiotics and grown at 37°C in 10 mL of LB. For T3 infection experiments cells were grown till OD_600_ of 0.6, when 200 μl aliquots were transferred to 96-well plates and infected at desired Multiplicity of Infection (MOI). To study the toxicity of SAMase expression, cells were grown till OD_600_ of 0.3 and induction with different concentrations of L-arabinose was done in 96-well plates. For the studies of SAMase expression on BREX defence, cells were grown till OD_600_ of 0.3, induced with the indicated concentration of L-arabinose and left with shaking for 30’ before 200 μl aliquots were transferred to 96-well plates and supplemented with phage at the indicated MOI. Optical density was monitored for 10-20 h. All experiments were performed in three biological replicas.

### Phage titer determination and efficiency of plating (EOP) assay

To determine the titer of active phage particles in cell lysates, the double agar overlay method was used. Overnight cultures of bacteria (100 *μ*l) were mixed with 10 ml of 0.6% top LB agar supplemented with appropriate antibiotics and poured on the surface of precast 1.2% bottom LB agar plates. 10 μl drops of serial 10-fold phage lysate dilutions were spotted on the top agar, allowed to dry and plates were incubated at 37°C overnight. The level of protection was determined as the ratio of phage titers obtained on a non-restrictive (BREX–) host relative to that on restrictive (BREX+) host. All experiments were performed in biological triplicates.

### One-step growth curve assay

Overnight culture of BL21 BREX+ or BREX-strains was diluted 100-fold in LB medium and cultivated in 10 ml at 37°C until OD_600_ = 0.6. At this point, phage was added to the culture at MOI=0.1. After this, cells were kept agitated 37°C and 1 ml aliquots were taken every 10 minutes. Cells were sedimented by centrifugation at 6,000g for 3 minutes. Phage titer in the supernatant was determined as described. After 12h of incubation at 30°C phage titer was calculated and the dynamics of phage production was estimated.

### Construction of T3Δ0.3 phage

Deletion of the *0*.*3* gene from T3 genome was achieved using λ RED-assisted SpCas9 genome editing.^37^ In short, a plasmid pKDsgRNA was designed to express a spacer against T3 *0*.*3* gene. pBAD vector for homologous recombination was provided that carried an insert with 300-bp 5’ and 3’ fragments bordering the region of desired deletion. pBAD and pKDsgRNA were transformed into cell carrying pSpCas9 vector. λ RED genes were expressed from the pKDsgRNA and the culture was infected with wild-type T3 in a range of MOIs in a 96-well plate format. SpCas9 counter-selection allows elimination of the wild-type phage, while λ RED system promotes homologous recombination between the phage genome and pBAD leading to removal of the *0*.*3* gene sequence, which generates escaper phages not-sensitive to CRISPR selection. Wells were screened by PCR for the presence of lysate with recombinant phages and T3*Δ0*.*3* was further purified from individual plaques and verified by Sanger sequencing (Evrogen).

### Pacific Bioscience sequencing

BREX, Dam and EcoKI modification frequencies were determined using the PacBio platform. Bacterial cultures were grown in triplicates in 10 ml of LB medium supplemented with appropriate antibiotic at 37°C for 1 h. Ocr or T3 SAMase expression was induced for an additional 2 h with 0.2% L-arabinose. Total DNA was purified from 2 ml of cultures using the GeneJET Genomic DNA Purification Kit (K0722, Thermo Scientific) in accordance with manufacturer’s instructions. The extracted DNA was sheared to a targeted mean size of 500 bp (the effective subreads size was on average of 959 bp (min: 828, median: 942, max: 1240)) using ultrasonicator (Covaris) and purified with AMPure PB beads (Pacific Biosciences). PacBio sequencing libraries were prepared using the SMRTbell Template Prep kit 1.0 (Pacific Biosciences). Protocols for polymerase binding were generated by the Pacific Biosciences Binding Calculator. Sequencing was performed using PacBio RS II (Pacific Biosciences), each sample being sequenced for six hours on one SMRT cell. The PacBio SMRT Analysis software was used for alignment of reads and detection of base modification. The percentage of modified BREX, EcoKI and Dam motifs was determined using the coverage and modification QV reported in the ‘modifications.csv.gz’ file produced by the analysis software. For each motif, methylation-targeted adenines with a reported coverage lower than 25 were discarded, and the remaining adenines were considered as modified if their modification QV was at least 30. Sequencing statistics are available in the Table S1.

### Study of the SAMase effects on λ phage genome modification

*E. coli* BW25113 cells were lysogenized with temperature-sensitive λ phage strain cI857 *bor*::*Cm*. The lysogen was transformed with pBTB-2/pBREX AL and pBAD T3 SAMase. The cultures were grown at 30°C to an OD_600_=0.6 in 10 ml LB medium supplemented with appropriate antibiotics in the presence of indicated L-arabinose concentrations to induce SAMase expression. λ prophage induction was achieved by incubating cells at 42°C for 60 min until lysis was observed, and cultures were then additionally treated with 100 *μ*l of chloroform. The lysate was cleared by centrifugation, and phage titers were determined on lawns of BREX-, BREX+ or AB1157 cells on LB plates supplemented with 0.2% maltose and 5 mM MgSO_4_. The plates were incubated overnight at 37°C and the level of protection was estimated as the ratio of phage titers obtained on non-restrictive (BREX-) host relative to restrictive (BREX+ or AB1157) hosts. The model of the experimental system allowing to produce modified phage λ progeny and estimate the effects of modification inhibition is presented on the Figure 4B.

### In vivo protein pull-down

Overnight cultures of *E. coli* strains with plasmids encoding the Strep-tagged protein (pBAD T3 SAMase C-strep or pTG MetK C-Strep) were diluted 100-fold in 500 ml of fresh LB medium supplemented with antibiotics and grown in 1 L flasks to OD_600_~0.3, induced with 0.2% L-arabinose and kept shaking until OD_600_ ~0.6. Cells were harvested by centrifugation at 4000g at 4°C for 10 min, cell pellets were washed in 10 ml of buffer StrepA (150 mM NaCl, 1 mM EDTA, 100 mM Tris–HCl; pH 8.0), and resuspended in the same buffer supplemented with a protease inhibitor cocktail (Roche). Cells were disrupted by sonication on ice (40% power, 10’’ pulse, 20’’ pause, 30 minutes on Qsonica sonicator with 6 mm sonotrode), and the lysate was clarified by centrifugation at 15 000g, 4°C for 30 min. The supernatant was applied to an equilibrated 1-ml StrepTrap HP chromatography column connected to the NGC Chromatography System (BioRad), washed with 20 CV of StrepA and elution was performed by applying a gradient concentration of StrepB (150 mM NaCl, 1 mM EDTA, 100 mM Tris–HCl, 2.5 mM desthiobiotin; pH 8.0). Protein-containing fractions were concentrated using 10-kDa Amicon centrifugal filter units (Merk) and analyzed by SDS-PAGE on a 4–20% gradient gel. The identity of protein bands was determined by matrix-assisted laser desorption/ionization time-of-flight (MALDI-TOF) mass spectrometry. Samples were prepared with Trypsin Gold (Promega) in accordance with manufacturer’s instructions. Mass spectra were obtained using the rapifleX system (Bruker).

### T3 SAMase expression and purification

TOP10 cells containing a pBAD T3 SAMase C-His vector encoding the C-terminally His6-tagged T3 SAMase (wild-type, E68Q/Q94A or other mutants) were inoculated in LB media containing 50 μg/ml ampicillin and incubated overnight at 37 °C with shaking (100 rpm). For protein expression, 5 ml overnight culture was used to inoculate 800 ml culture in the same media. The flasks were incubated at 37 °C until OD_600_ reached 0.4, when expression was induced with 0.2% L-arabinose. The flasks were further incubated at 37 °C for 6 h and 18 °C overnight. The cells were harvested by centrifugation and the pellet was resuspended in lysis buffer (50 mM Tris-HCl pH 8, 300 mM NaCl, 20 mM imidazole, 5 mM β-ME) with complete EDTA-free protease inhibitor (Roche). The cells were lysed by sonication (Vibra-Cell™ VCX 130) on ice with 80% amplitude in 2 second pulses and 8 second pauses for 25 min. The lysate was centrifuged 45 min at 4 °C and 39 000 g. The supernatant was filtered through a 0.45-μm-syringe filter, loaded to a gravity column containing Ni-Sepharose (GE Healthcare) pre-equilibrated with buffer A (50 mM Tris-HCl pH 8, 300 mM NaCl, 40 mM imidazole, 5 mM β-ME) and incubated under slow rotation for 30 min at 4 °C. The column was washed extensively with buffer A and the protein was eluted with elution buffer (50 mM Tris-HCl pH 8, 300 mM NaCl, 500 mM imidazole, 5 mM β-ME). Fractions containing the protein were loaded onto a HiLoad 16/60 Superdex 200 column equilibrated with buffer B (50 mM Tris-HCl pH 8, 300 mM NaCl, 5 mM β-ME). Peak fractions corresponding to the T3 SAMase-MetK complex were used directly for cryo-EM or pooled and concentrated to 5-10 mg/ml for SAXS experiments. Gel filtration of the wild-type and T4A SAMase:MetK complexes eluted after His-tag purification was performed on a Superdex 200 Increase 10/300 column (Cytiva) in StrepA buffer at 0.5 ml/min flow rate.

### In vitro SAM cleavage reaction

Wild-type or mutant C-terminally His-tagged T3 SAMase for the *in vitro* SAM cleavage studies was obtained in a similar manner. Protein was produced from 300-600 ml of LB and purified on a 1ml HisTrap column connected to the NGC Chromatography System (BioRad). After lysate loading column was washed with 10CV of buffer A, followed by 4CV gradient elution with buffer B. Buffer was exchanged in protein containing fractions through overnight dialysis against 1L of StrepA (150 mM NaCl, 1 mM EDTA, 100 mM Tris–HCl; pH 8.0), protein concentration was measured with a Qubit protein assay kit (Invitrogen) and Quick Start Bradford Dye (Bio-Rad). 500 nM SAMase was incubated with 1 mM SAM (Heptral) in 30 μl in the following buffer: TrisHCl, pH 7 – 100 mM; KCl – 100 mM. The reaction was carried at room temperature and at indicated time points was stopped by the addition of 2V of 50 mM citric acid (pH=2.9) and aliquots were kept frozen at -80°C. A control reaction was carried with SAM without the addition of SAMase, to check spontaneous SAM decay, and with SAMase without the addition of SAM, to check for the absence of MTA traces in the protein sample.

SAM (substrate) and MTA (one of the products) of the *in vitro* SAM cleavage reaction were separated by reverse-phase high performance liquid chromatography (HPLC) with Zorbax Eclipse Plus C^18^ 5-μm (4.6 by 250 mm) column (Agilent) connected to the 1220 Infinity II LC system (Agilent). The column was equilibrated in 0.1% TFA for 3 minutes and then compounds were separated in a linear gradient of acetonitrile (0 to 16%) for 10 minutes, followed by 5 minutes wash with 80% acetonitrile. The identity of compounds was verified by comparing the retention time of peaks observed in an experimental sample compared to those for the MTA and SAM standards.

### Cryo-EM grid preparation and data collection

Wild-type T3 SAMase-MetK complex directly after size-exclusion chromatography was mixed with an equal volume of water to adjust the buffer to 25 mM Tris-HCl pH 8, 150 mM NaCl, 2.5 mM β-ME and slightly concentrated for grid preparation. The cryo-EM specimens were prepared on C-Flat 1.2/1.3 holey carbon support on 300 mesh copper grids (Protochip, USA) that were glow-discharged using a PELCO EasiGlow glow discharge system (Ted Pella Inc, USA). 3 μl of 0.125 mg/ml complex was loaded onto the pretreated grid and blotted for 3.0 seconds before plunge-freezing in liquid nitrogen-cooled ethane, using a Vitrobot Mark-IV (ThermoFisher Scientific, USA).

Data collection was performed at SciLifeLab in Solna, Sweden, with a Titan Krios G3 microscope (ThermoFisher Scientific, USA) equipped with a K3 camera (Gatan K3 Bioquantum, Inc) and energy filter (20 eV slit width). Micrographs were recorded at 105,000x magnification, equivalent to a nominal pixel size of 0.8617 Å in the imaging plane. The electron dose of 47.75 e/Å^2^ was fractionated into 40 frames and the average exposure time per image was 3.0 seconds. Movies were collected using the EPU software (v2.11). Images were CTF-corrected by phase flipping and amplitude correction. Data-collection parameters are summarized in Table S2.

### Cryo-EM data processing

Data was processed by using CryoSPARC v3.3.2+.^63^ From 5,447 movies, 3,747 micrographs were accepted. Particles were automatically picked by the cryoSPARC Blob picker and subjected to 2D classification. Template picking resulted in 1,445,186 particles that were subjected to several rounds of 2D classification. Homogeneous refinement of 540,185 particles applying C1 symmetry produced an initial map of 3.14 Å resolution. After CTF refinement and 3D classification, 525,534 particles were subjected to heterogeneous refinement applying C1 symmetry. Particles with full occupancy of T3 SAMase were subjected to homogeneous refinement with D2 symmetry, resulting in a 3.0 Å map. Non-uniform refinement improved the resolution to 2.8 Å. Symmetry expansion and local refinement with C1 symmetry using a mask around the SAMase produced a 3.0 Å map of the T3 SAMase. The pixel size was calibrated against PDB 1P7L^64^ by rigid-body fitting and scaling of the voxel size in ChimeraX^65^ to be 0.8343 Å. The data processing scheme is summarized in Figure S4.

### Model building and refinement

Model building of T3 SAMase in the map from local refinement was assisted by initial models from RaptorX^66^ and AlphaFold2.^67^ The model was manually rebuilt in Coot (v. 0.9.3),^68^ and real-space refined in Phenix (v. 1.20.1-4487)^69^ using reference model restraints and Ramachandran restraints. The MetK model was built by rigid-body fitting and manual rebuilding of MetK based on PDB entry 1P7L^64^ (into the non-uniform refined map, followed by real-space refinement in Phenix. The complete MetK-T3 SAMase complex was real-space refined in the map from non-uniform refinement with Ramachandran restraints, and water molecules were assigned automatically by Coot and individually checked. Structure figures were prepared with ChimeraX^65^ and PyMol.^70^

### SAXS

SEC-SAXS data for the T3 SAMase-MetK complex were collected at the Diamond Light Source on beamline B21. In-line SEC-SAXS was performed using an Agilent 1200 HPLC system connected to a Shodex KW403 column. Wild-type T3 SAMase-MetK complex at 4.9 mg/ml and T3 SAMase E68Q/Q94A-MetK complex at 9.6 mg/ml were loaded onto the column equilibrated with 50 mM Tris-HCl pH 7.5, 300 mM NaCl, 5 mM β-ME and 2 % glycerol. The data was initially subtracted with the buffer and data processing was performed using ScÅtter.^71^ Further data analysis was performed with Primus^72^ and the FOXS server.^73^

### Mass photometry

Molecular weight determination of a 30 nM wild-type T3 SAMase-MetK complex was performed by mass photometry using a Refeyn One^MP^ instrument. The mass was determined by the highest count number.

### Western blot

To estimate *in vivo* levels of T3 SAMase wild-type and mutated variants a western-blot with His-tag specific antibodies was applied. As an internal reference we estimated the signal form the alpha subunit of the host RNA-Polymerase. Bacterial cultures were grown in 20 mL LB at 37°C until OD_600_=0.2, induced with 0.2% L-arabinose, kept agitated for 2.5 hours for SAMase production and then placed on ice for 10 minutes. Cultures were equilibrated to OD_600_ = 0.65 in 20 mL, collected by centrifugation at 4°C, 3600g for 13 minutes and resuspended in 1 mL of PBS supplemented with 1mM PMSF. Lysates were obtained by sonication on ice (20% power, 5’’ pulse, 5’’ pause, 4.5 minutes on Qsonica sonicator with 2 mm sonotrode) and clarified for 1 hour at 4°C, 21000g. 2 identical gels were prepared for the staining of SAMase or refence protein. 15 μl of lysate were run in 16% SDS-PAGE followed by transfer on the PVDF membrane using TransBlot Turbo device (Bio-Rad). Membrane was washed in mQ and incubated at +4°C overnight in 10 mL of PBS with 5% non-fat dry milk. 1-2 μl of primary anti-His (PA1-983B, Thermo Scientific) or anti-alpha RNAP (4RA2, Neoclone) antibodies were added and Tween20 was supplied to 0.05%, incubation was carried for 2 hours at room temperature, followed by 3 washes in 20 ml 0.05% Tween20/PBS. 1 μl of secondary antibodies (ab205719 or ab6721) were added in 12 ml of 0.05% Tween20/PBS/1% BSA and incubation was carried at room temperature for 1 hour, followed by 3 washes. Immunodetection was carried with Clarity Western ECL Substrate (Bio-Rad). Images were received on a ChemiDoc XRS+ system (Bio-Rad) and bands intensity was calculated in ImageJ.

### SAMase Toxicity Assay

SAMase toxicity was estimated either in a liquid culture, as described above, either in a spot-test assay. Cells were grown in 10 ml of LB and equilibrated to OD=0.6 followed by serial 10-fold dilution droplets plating on LBA supplemented with required antibiotics and optional inducer.

## Data availability

Atomic coordinates for the T3 SAMase-MetK structure have been deposited in the PDB with accession code 8BB1. Maps have been deposited in the Electron Microscopy Data Bank with accession codes EMD-15953 (full complex) and EMD-15952 (local map for T3 SAMase). SAXS data have been deposited in the SASDB with accession codes SASDQ95 (complex of wild type T3 SAMase with MetK) and SASDQA5 (complex of T3 SAMase E68Q/Q94A with MetK).

## Acknowledgements

The phage T3 was provided by the Cristian Aparicio-Maldonado and Dr. Franklin Nobrega, while Dr Shimon Bershtein kindly shared with us *E. coli* strains with *metK* gene substitutions. We thank Dr Daniel Larsson, CryoScreeNet, for cryo-EM data collection and for help and advice regarding data processing. We are grateful for access to beamline B21 at the Diamond light source, Didcot, UK (measurements in May and June 2021).

---

**Figure S1.**
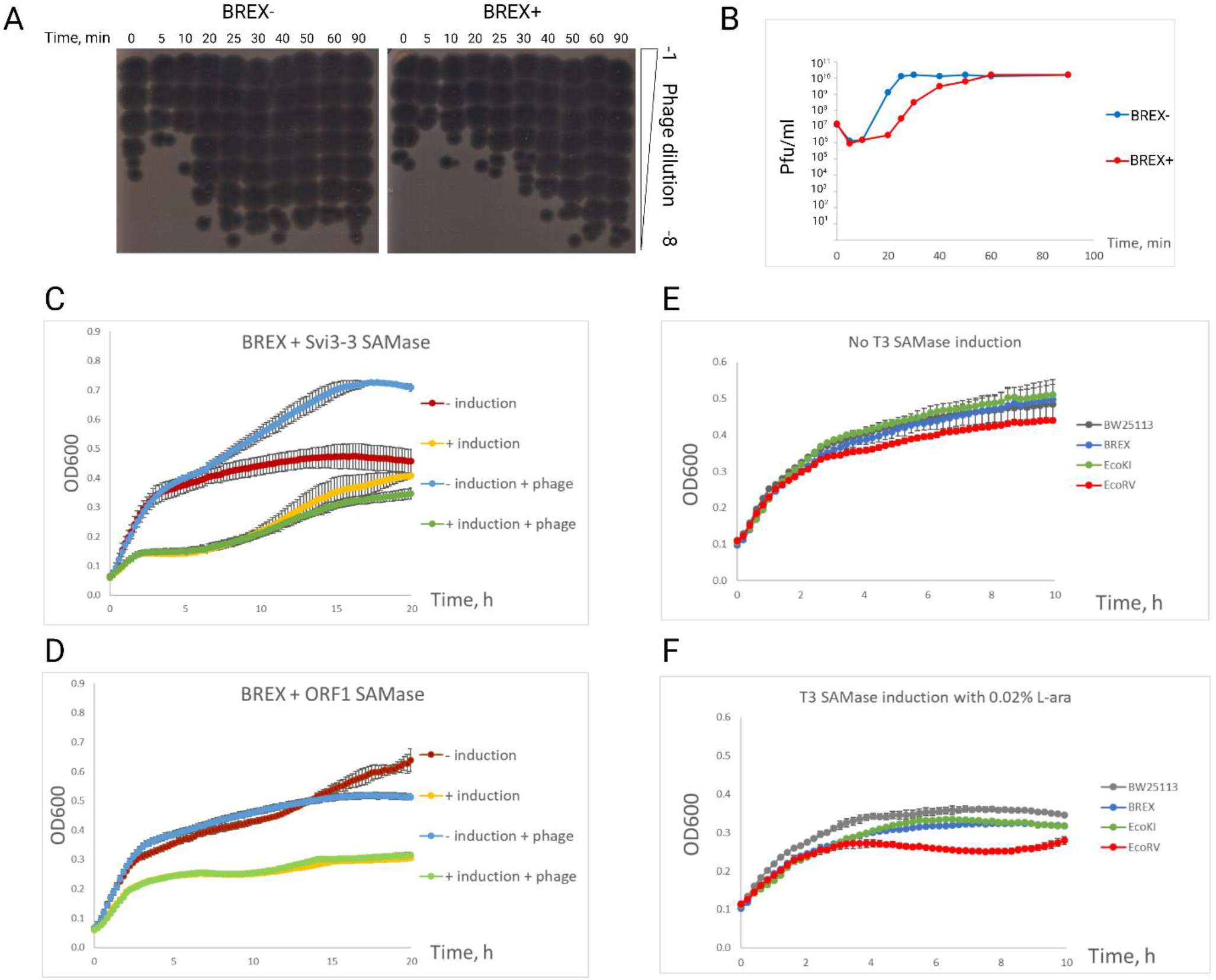
(A)One-step growth curves assay with phage T3 and BL-21 BREX- or BREX+ culture. Phage was added at *t=0* at MOI=0.1 and aliquots were taken at indicated time points to measure the titer of free phage particles on the BREX-lawn. (B)Corresponding phage titers for the experiment shown in (A). Presented are results for one out of two conducted independent experiments. (C-D) Expression of T3 SAMase homologs does not provide an anti-BREX effect. T7*Δ0*.*3* infection of BW25113 BREX+ culture expressing different SAMase variants. Cultures were supplemented with 1mM IPTG to induce SAMase expression from pCA24N (Svi3-3) (C) or pCA24N (ORF1) (D) vectors. MOI=0.001, phage was added at *t=0*. (E-F) SAMase expression increases toxicity in the strain bearing Type II R-M system EcoRV, but not EcoKI or BREX defence systems, compared to BW25113 control. Growth of cultures without induction (E) or with 0.02% L-arabinose induction (F) of SAMase expression in BW25113, BW25113 + pBREX AL, BW25113 + pBR322 EcoKI, or BW25113 + pEF42 EcoRV.

**Figure S2.**
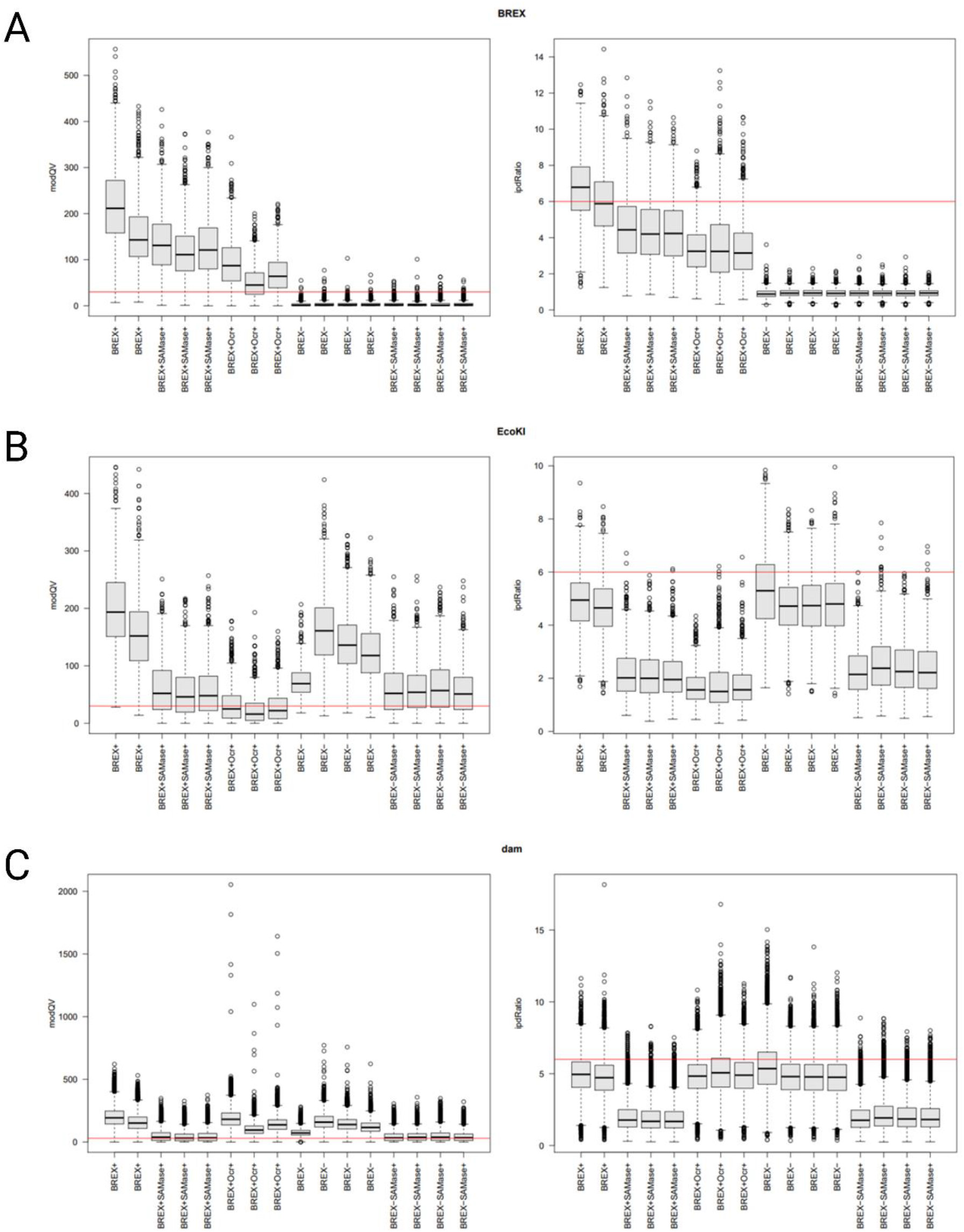
(A-C) Effects of T3 SAMase or Ocr on adenine-specific methylation in the *E. coli* genome. Distribution of the modification QVs and of the IPD ratios for BREX (A), EcoKI (B), and Dam (C) sites in the genomes of BW25113 (EcoKI *m+r-*) BREX- or BREX+ cells expressing, where indicated, Ocr or T3 SAMase from pBAD vector, as revealed by Pacific Bioscience sequencing. Data obtained from independent replicates are shown. Red lines label a modification QV of 30 and an IPD ratio of 6, which indicates a six-fold longer than expected adenine incorporation time.

**Figure S3.**
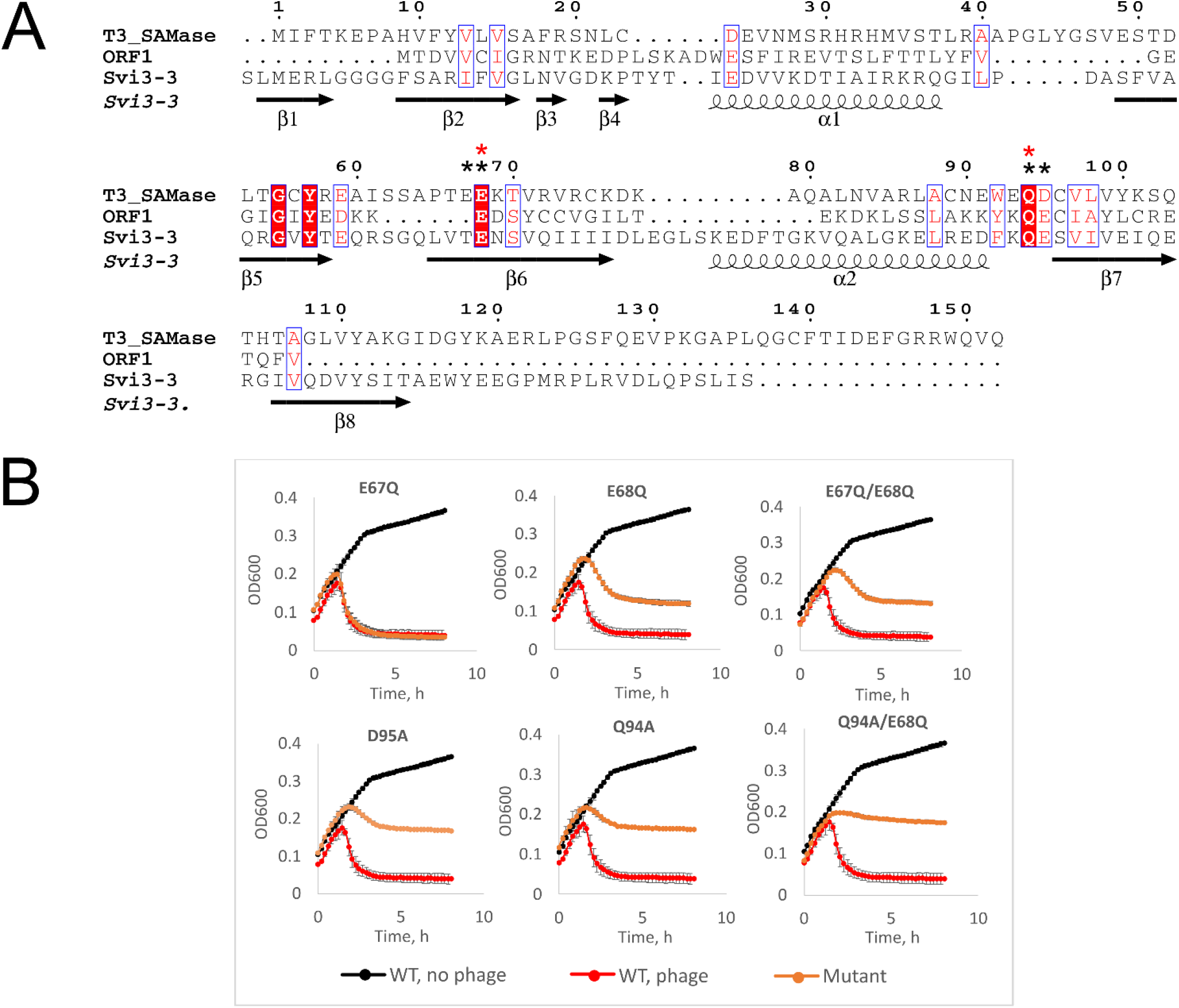
(A)Multiple-sequence alignment of T3 SAMase with other phage-encoded SAMases.^1^ Strictly conserved positions are shown in white on red background and conservatively substituted positions in red on white background. Secondary structure of Svi3-3 (PDB 6ZM9) is indicated above the alignment. Black stars above the alignment indicate positions of single-substitution mutants, red stars denote the catalytically-deficient double mutant used throughout the work. The figure was prepared using ESPript 3.0.^2^ (B)Mutations in T3 SAMase decrease but do not completely eliminate anti-BREX activity. Growth curves of BREX+ culture infected with phage T7*Δ0*.*3* at MOI = 0.1 in the presence of different C-His SAMase variants induced with 0.2% arabinose. Phage was added at *t=0*.

**Figure S4.**
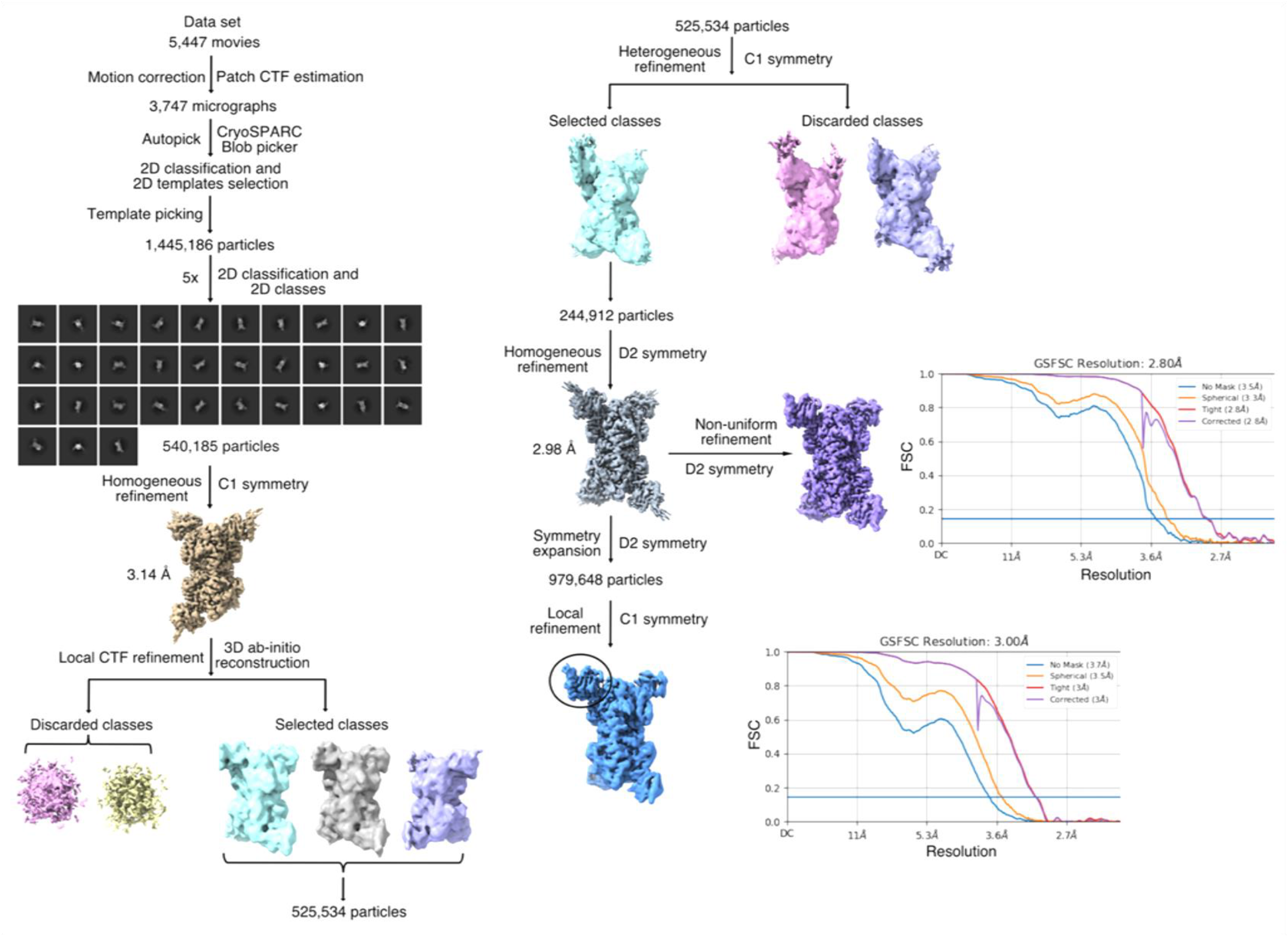
Cryo-EM structure determination of the T3 SAMase-MetK complex.

**Figure S5.**
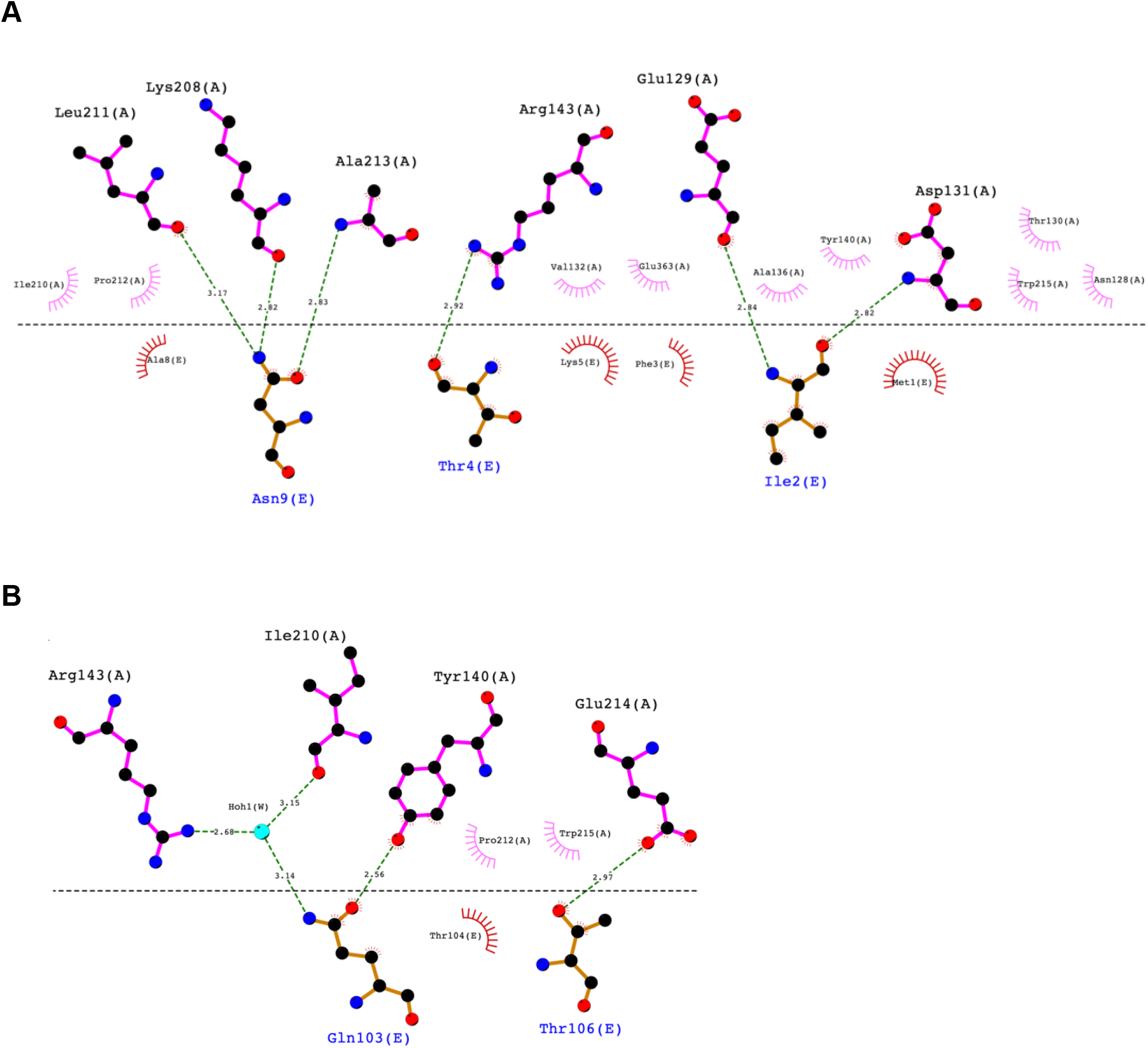
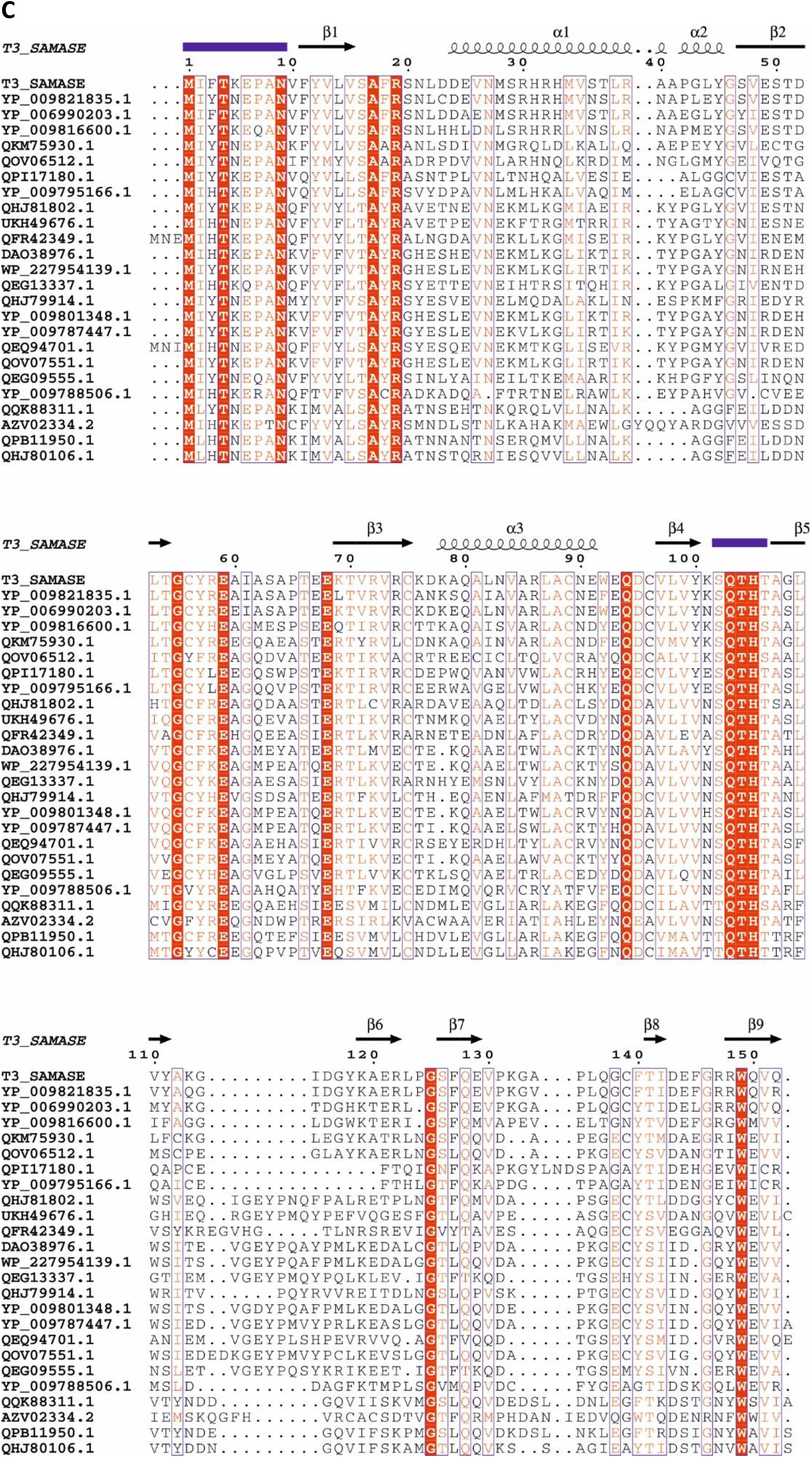
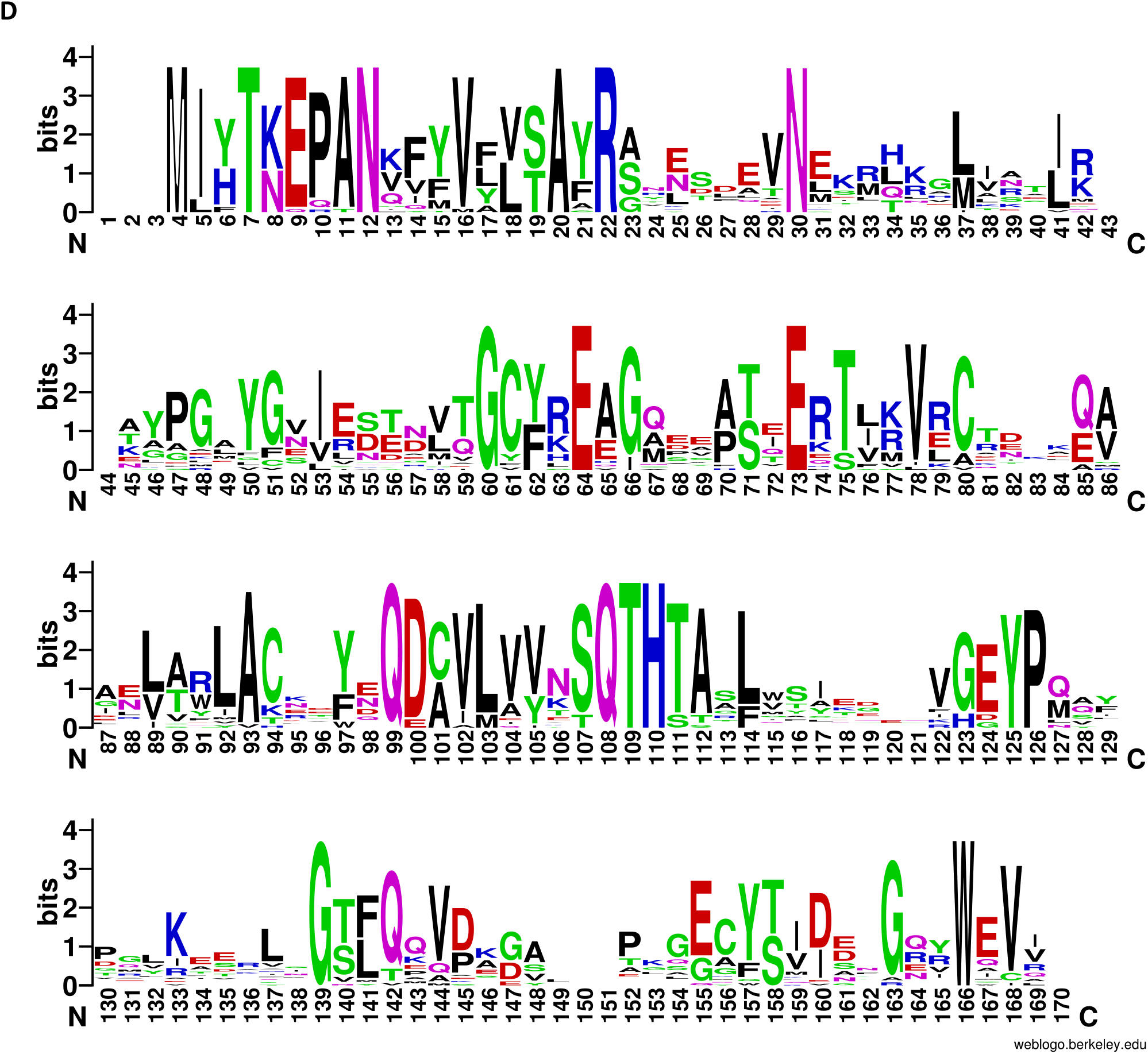
(A-B) Detailed interactions between T3 SAMase and MetK. (A) Interactions of the N-terminal tail of T3 SAMase with MetK. (B) Interactions of the β4-β5 loop of T3 SAMase with MetK. The figure was prepared with LigPlot+ v.2.2.5.^3^ (C-D) Sequence comparison of T3 SAMase with representative similar SAMase sequences. (C) Multiple sequence alignment of representative sequences of at least 30% sequence identity and 90% coverage to T3 SAMase from the clustered non-redundant NCBI database. Secondary structure of T3 SAMase is indicated above the alignment. The N-terminal tail and the β4-β5 loop are indicated with blue bars. Figure was prepared using ESPript.^2^ (D) Sequence logo of aligned representative sequences shown in C. Out of the residues interacting with MetK, T4 (7 in the logo), Q103, T104 and H105 (108-110 in the logo, H105 interacts with the N-terminal tail that in turn interacts with MetK) are strictly conserved. Figure was prepared with WebLogo.^4^

**Figure S6.**
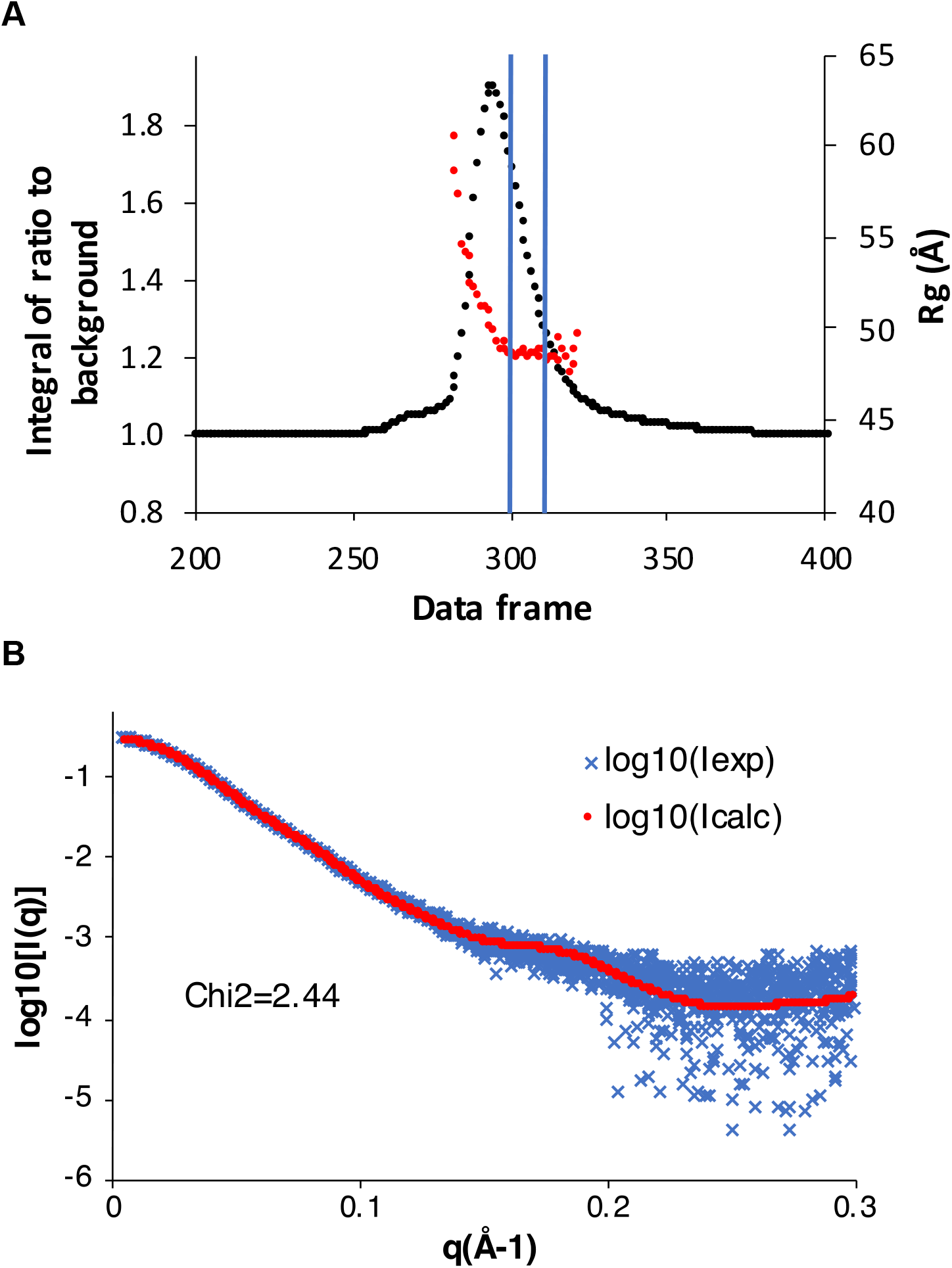
(A)SEC-SAXS scattering intensity plot for the E68A/Q94A T3 SAMase-MetK complex. Red markers indicate the radii of gyration (R_g_) calculated from the individual scattering curves. Blue bars indicate the data frames used for further analysis. (B)Overlay of experimental scattering data (blue) and the calculated scattering curve based on the 4:4 wild-type T3 SAMase-MetK cryo-EM complex structure (red).

**Figure S7.**
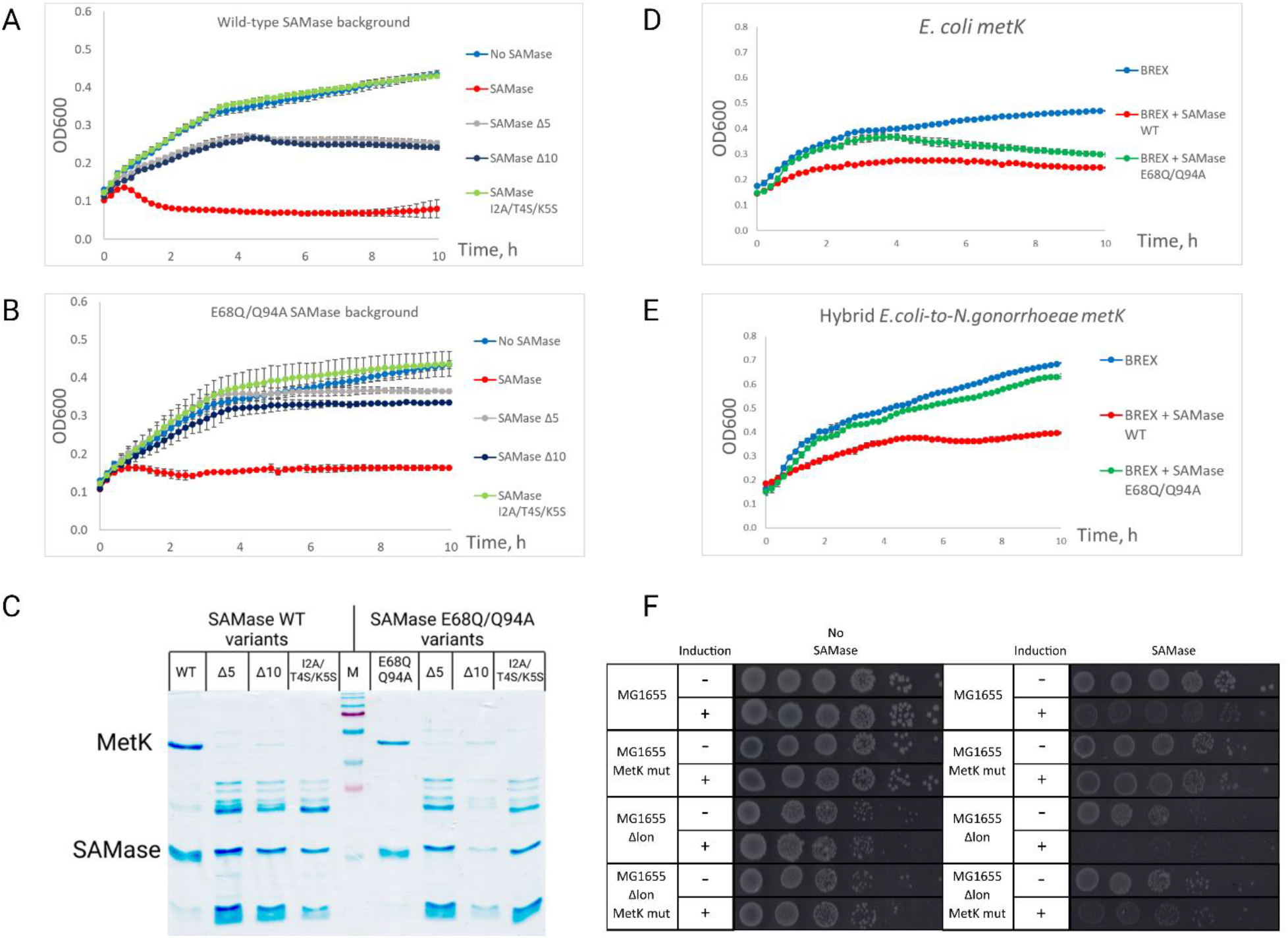
(A-B) Growth curves of BREX+ culture carrying C-His SAMase wild-type (A) or E68Q/Q94A (B) with additional N-terminal mutations and infected with phage T7*Δ0*.*3* at MOI = 0.01, induction with 0.2% arabinose. Phage was added at *t=0*. (C)16% SDS-PAGE showing the results of pull-downs from lysates of *E. coli* BW25113 expressing His-tagged SAMase from pBAD vector (wild-type or E68Q/Q94A with additional mutations). Δ5, Δ10 and I2A/T4S/K5S variants were concentrated with Amicon centrifugal filter units, which results in stronger background of contaminant protein bands. The identity of all SAMase and MetK bands was confirmed using MALDI-TOF MS analysis. M – Marker, PageRuler Prestained Plus. (D-E) Growth curves of the MG1655 wild-type (D) or *E*.*coli-to-N*.*gonorrhoeae* MetK (E) variants expressing SAMase wild-type or E68Q/Q94A mutants in the presence of BREX. SAMase expression was induced with 0.2% L-arabinose, 30 minutes prior to *t=0* point. (F) Toxicity assay with MG1655 cells expressing SAMase wild-type or E68Q/Q94A in *Lon*^*+*^*/Lon*^*-*^ and MetK WT/*E*.*coli-to-N*.*gonorrhoeae* backgrounds. Induction with 0.02% L-arabinose. Wild-type SAMase demonstrates enhanced toxicity in cells lacking Lon protease, while disruption of SAMase:MetK complex (*E*.*coli-to-N*.*gonorrhoeae* MetK / *Lon*^*-*^ background) partially restores viability.

