## Supplementary material for "Phage T3 overcomes the BREX defence through SAM cleavage and inhibition of SAM synthesis": Table S2

**Supplementary table 2**

Data collection, refinement parameters and model validation

| **Data collection** | | |
| --- | --- | --- |
| Microscope model | TFS Krios | |
| Magnification | 105,000x | |
| Electron dose | 47.75 e/Å^2^ | |
| Acceleration voltage (kV) | 300 | |
| C2 aperture diameter (µm) | 50.0 | |
| Detector | Gatan K3 Bioquantum (6k x 4k) | |
| Exposure time (seconds) | 3.0 | |
| Target defocuses | -2.6 µm to -0.6 µm in 0.2 µm steps | |
| Number of frames | 40 | |
| Dose rate | 11.818 e/px/s | |
| Number of grids imaged | 1 | |
| Number of movies | 5,447 | |
| Number of accepted micrographs | 3,747 | |
| Number of particles selected from the micrographs | 1,445,186 | |
| **3D reconstruction** | | |
| Model | MetK | T3 SAMase |
| Resolution | 2.8 Å | 3.0 Å |
| Number of particles used for final reconstruction | 244,912 | 979,648 |
| Refinement method | Non-uniform refinement | Local refinement |
| Applied point symmetry | D2 | C1 |
| **Model refinement and validation** | | |
| Refinement package | Phenix v. 1.20.1-4487 | |
| Refinement space | Real | |
| **Model statistics** | | |
| Chains | 8 | |
| Total atoms | 16,168 | |
| Protein residues | 2,100 | |
| Water | 24 | |
| Cl^-^ ions | 4 | |
| MolProbity score | 1.16 | |
| Clash score | 3.65 | |
| RMSD bond length (Å) | 0.003 | |
| RMSD bond angles (°) | 0.415 | |
| Ramachandran (%) (Favored/Allowed/Outlier) | 97.97 / 2.03 / 0.00 | |
| Rama-Z (whole/helix/sheet/loop) | 0.54 / 1.25 / 0.54 / -0.29 | |
| Rotamer outliers (%) | 0.71 | |
| Cβ outliers (%) | 0.00 | |
| CaBLAM outliers (%) | 0.79 | |
