## Supplementary material for "Phage T3 overcomes the BREX defence through SAM cleavage and inhibition of SAM synthesis": Table S3

**Supplementary Table S3. Bacterial strains, plasmids, and phages used in the study.**

| ***E. coli* strain** | **Comments** | **Source** |
| --- | --- | --- |
| BW25113 | *E. coli* K12 F^–^ *Δ*(*araD-araB*)567 *ΔlacZ*4787(::*rrnB*-3) λ– *rph*-1 *Δ*(*rhaD-rhaB*)568 *hsdR*514 | Lab stock |
| BL21(DE3) | *E. coli* B F– *omp*T *hsdS_B_* (r_B_–, m_B_–) *gal dcm* (DE3) | Lab stock |
| AB1157 | *E. coli* K12 F^–^ *thr-*1 *araC*14 *leuB*6(Am) *Δ(gpt-proA)*62 *lacY*1 *tsx-*33 *qsr'-*0 *glnX*44(AS) *galK*2(Oc) λ^-^ *Rac-*0 *hisG*4(Oc) *rfbC*1 *mgl-*51 *rpoS*396(Am) *rpsL*31 *kdgK*51 *xylA*, *mtl-*1 *argE*3(Oc) *thiE*1, Str^R^ | Lab stock |
| XL1-Blue | *E. coli* K12 *recA*1 *endA*1 *gyrA*96 *thi*-1 *hsdR*17 *supE*44 *relA*1 *lac* [*F'proAB lacI^q^ZΔM*15 *Tn*10], Tet^R^ | Evrogen |
| TOP10 | *E. coli* K12 F- *mcrA (mrr-hsdRMS-mcrBC) 80lacZ M15 lacX74 recA1 ara139 (ara-leu) 7697 galU galK rpsL endA1 nupG*, Str^R^ | Thermo Scientific |
| BW25113 λ_ts_ lysogen | λ *cI*_857_ *bor::Cm,* Cm^R^ | 1 |
| BW25113 *ΔhsdM* | KEIO collection, *hsdM::Kn*, Kn^R^ | 2 |
| BW25113 *Δlon* | KEIO collection, *lon::Kn*, Kn^R^ | 2 |
| BW25113 *ΔclpX* | KEIO collection, *clpX::Kn*, Kn^R^ | 2 |
| MG1655 | *E. coli* K12 F-, λ-*, ilvG-, rfb-50, rph-1* | Lab stock |
| MG1655 *Δlon* | P1 transduction from BW25113 *Δlon* | This work |
| MG1655 *ΔclpX* | P1 transduction from BW25113 *ΔclpX* | This work |
| MG1655 *E.coli-to-Ureaplasma urealyticum MetK* | MG1655 *metK::metK from U. urealyticum* + pFlag *U. urealyticum metK*, Amp^R^ | Dr. Shimon Bershtein |
| MG1655 *E.coli-to-Neisseria gonorrhoeae* *MetK* | MG1655 *metK* (D132P, V133T, I178V, I211V, A214P, W26L) | Dr. Shimon Bershtein |
| **Phage** | **Comments** | **Source** |
| T3 |  | Dr. Stan Brouns |
| T3 *Δ0.3* | *0.3* gene scarless deletion | 3 |
| T7 |  | Lab stock |
| T7 *Δ0.3* | *0.3*::*trxA* | 4 |
| λ_vir_ | λ mutant with an obligatory lytic lifecycle | Lab stock |
| **Plasmids** | **Comments** | **Source** |
| pBAD L24 | pBAD/His B *Δ312-455* (lacks features for protein purification) with introduced EcoRI and SacI restriction sites, Amp^R^ | Lab stock |
| pBAD T7 Ocr | pBAD L24 with T7 Ocr, araBAD promoter, Amp^R^ | 4 |
| pBAD T3 SAMase | pBAD L24 with 152 amino-acid long T3 SAMase product, araBAD promoter, Amp^R^ | This work |
| pBAD T3 SAMase short | pBAD L24 with 125 amino-acid long T3 SAMase product, araBAD promoter, Amp^R^ | This work |
| pBAD T3 SAMase C-His | pBAD T3 SAMase encoding C-terminally His_6_-tagged SAMase variant, Amp^R^ | This work |
| pBAD T3 SAMase C-Strep | pBAD T3 SAMase encoding C-terminally Strep-tagged SAMase variant, Amp^R^ | This work |
| pBAD T3 SAMase X | A series of T3 SAMase mutants, where X denotes mutation - E67Q, E68Q, Q94A, D95A, E67Q/E68Q, E68Q/Q94A, Δ10, Δ5, I2A/T4S/K5S, ΔI2, T4A, Q103A, T104A | This work |
| pCA24N Orf1 | Viral SAMase identified in metagenomic DNA library, *lac* promoter, Cm^R^ | 5 |
| pCA24N Svi3-3 | Viral SAMase identified in metagenomic DNA library, *lac* promoter, Cm^R^ | 5 |
| pBTB-2 | Kn^R^ | 1 |
| pBREX AL | 6 genes BREX cluster from *E. coli* HS in low copy number vector pBTB-2, Kn^R^ | 1 |
| pCA24N MetK | ASKA collection, *lac* promoter, Cm^R^ | 6 |
| pTG | Cm^R^ | 1 |
| pTG MetK C-Strep | pTG encoding C-terminally Strep-tagged MetK variant, *araBAD* promoter, Cm^R^ | This work |
