## Supplementary material for "Phage T3 overcomes the BREX defence through SAM cleavage and inhibition of SAM synthesis": Table S4

**Supplementary Table S4. Primers used in the study.**

| **Name** | **Sequence (5′->3′)** | **Purpose** |
| --- | --- | --- |
| pBAD_F | ATGCCATAGCATTTTTATCC | Verification of cloning in pBAD |
| pBAD_R | GATTTAATCTGTATCAGGCTG |  |
| T3_SAMase_F | TTTGGGCTAACAGGAGGAAGAATTCATGATTTTCACTAAAGAGCC | Cloning of T3 SAMase |
| T3_SAMase_R | GCCAAGCTTGCGGCCGCGAGCTCTCAATGTTAATCATAAAGGCCA |  |
| T3_SAMase_short_F | TTTGGGCTAACAGGAGGAAGAATTCATGAGCAGACACCGC |  |
| T3_SAMase_Strep_R | CAGCCAAGCTTGCGGCCGCGAGCTCTTATTTTTCGAACTGCGGGTGGCTCCATTGTACTTGCCAGCG | Addition of tag to T3 SAMase |
| T3_SAMase_His_R | CAGCCAAGCTTGCGGCCGCGAGCTCTTAATGATGATGATGATGATGTTGTACTTGCCAGCG |  |
| MetK_Strep_F | TGTTTCTCCATACCCGTTTTTTGGGCTAACAGGAGGAGAATTCATGGCAAAACACCTTTTTAC | Cloning of E. coli *metK* with Strep-tag |
| MetK_Strep_R | CCGCCAAAACAGCCAAGCTTGCTTATTTTTCGAACTGCGGGTGGCTCCACTTCAGACCGGCAGCATC |  |
| T3_D95A_F | TGGGAGCAAGCTTGCGTACTG | T3 SAMase mutagenesis |
| T3_D95A_R | CTCATTACAAGCTAGGCG |  |
| T3_Q94A_F | TGAGTGGGAGGCAGATTGCGTAC |  |
| T3_Q94A_R | TTACAAGCTAGGCGTGCA |  |
| T3_E67Q_F | CGCACCAACTCAGGAAAAAAC |  |
| T3_E67Q_R | CTTGAGATTGCCTCACGATAG |  |
| T3_E68Q_F | ACCAACTGAGCAGAAAACTGTTCGTGTAC |  |
| T3_E68Q_R | GCGCTTGAGATTGCCTCA |  |
| T3_E67Q-E68Q_F | CGCACCAACTCAGCAGAAAACTGTTCG |  |
| T3_E67Q_R | CTTGAGATTGCCTCACGATAG |  |
| T3_D95A_F | TGGGAGCAAGCTTGCGTACTG |  |
| T3_SAM_dN5_F | TTTGGGCTAACAGGAGGAAGAATTCATGGAGCCTGCGAAC |  |
| T3_SAM_dN10_F | TTTGGGCTAACAGGAGGAAGAATTCATGTTCTATGTACTGGTTTC |  |
| SAMase_Q103A_F | ATACAAATCAGCGACTCACACGGCTGG |  |
| SAMase_Q103A_R | ACCAGTACGCAATCTTGC |  |
| SAMase_T104A_F | CAAATCACAGGCGCACACGGCTG |  |
| SAMase_T104A_R | TATACCAGTACGCAATCTTG |  |
| SAMase_T4A_F | CATGATTTTCGCGAAAGAGCCTG |  |
| SAMase_T4A_R | AATTCTTCCTCCTGTTAG |  |
| SAMase_I2del_F | TTCACTAAAGAGCCTGCG |  |
| SAMase_I2del_R | CATGAATTCTTCCTCCTGTTAG |  |
